# A Proximity Biotinylation Assay with a Host Protein Bait Reveals Multiple Factors Modulating Enterovirus Replication

**DOI:** 10.1101/2022.05.24.493328

**Authors:** Seyedehmahsa Moghimi, Ekaterina Viktorova, Samuel Gabaglio, Anna Zimina, Bogdan Budnik, Bridge G. Wynn, Elizabeth Sztul, George A. Belov

## Abstract

As ultimate parasites, viruses depend on host factors for every step of their life cycle. On the other hand, cells evolved multiple mechanisms of detecting and interfering with viral replication. Yet, our understanding of the complex ensembles of pro- and anti-viral factors is very limited in virtually every virus-cell system. Here we investigated the proteins recruited to the replication organelles of poliovirus, a representative of the genus *Enterovirus* of the *Picornaviridae* family. We took advantage of a strict dependence of enterovirus replication on a host protein GBF1, and established a stable cell line expressing a truncated GBF1 fused to APEX2 peroxidase that effectively supported viral replication upon inhibition of the endogenous GBF1. This construct biotinylated multiple host and viral proteins on the replication organelles. Among the viral proteins, the polyprotein cleavage intermediates were overrepresented, arguing that the GBF1 environment is linked to the viral polyprotein processing. The proteomics characterization of biotinylated host proteins identified those previously associated with the enterovirus replication, as well as more than 200 new factors recruited to the replication organelles. RNA metabolism proteins many of which normally localize in the nucleus constituted the largest group, underscoring the massive release of nuclear factors in the cytoplasm of infected cells and their involvement in the viral replication. Analysis of several newly identified proteins revealed both pro- and anti-viral factors, including a novel component of infection-induced stress granules. Depletion of these proteins similarly affected the replication of diverse enteroviruses indicating broad conservation of the replication mechanisms. Thus, our data significantly increase the knowledge about the organization of enterovirus replication organelles and may provide new targets for anti-viral interventions.

---

Enterovirus infections are associated with numerous life-threatening and/or economically important diseases ranging from the common cold to fatal encephalitis. Among multiple pathogenic enteroviruses, licensed vaccines are available only against poliovirus and Enterovirus A71, and there are no drugs approved by the FDA to control enterovirus infections (1-5). This appalling scarcity of effective anti-enterovirus measures visibly reflects the inadequate understanding of the molecular mechanisms underlying the development of infection of this important group of viruses.

The *Enterovirus* genus belongs to the family *Picornaviridae* encompassing small positive-strand RNA viruses with non-enveloped icosahedral capsids infecting vertebrate hosts. Poliovirus is the best-studied enterovirus, its single genome RNA of about 7500 nucleotides is polyadenylated on the 3’-end and has a small protein Vpg covalently attached to the 5’-end. The 5’-end long non-translated region of the genome RNA contains an internal ribosome entry site (IRES), and both 5’ and 3’ non-translated regions, as well as the coding part of the genome, contain cis-acting elements important for the RNA replication (6). The viral RNA is translated into a single polyprotein of about 200KDa which is processed by three viral proteases into three capsid and ten replication proteins, including stable intermediate products of the polyprotein processing (7-10). Upon accumulation of the replication proteins, they form a replication complex where most of the viral proteins are assembled *in cis*, i. e. processed from the same polyprotein precursor, and likely initiate replication of the same RNA that served as an mRNA for protein synthesis (11-14). The newly synthesized genomes can enter subsequent translation-replication cycles or be packaged into new viral particles.

The small genome size and consequently a limited repertoire of viral proteins implies that multiple host factors should support the replication process. Over the years, many host proteins that are required for, or facilitate the development of enterovirus infection have been identified (many of them are reviewed in (15-17), but the full catalog of all the cellular proteins that have been implicated in the enterovirus life cycle is yet to be compiled). The two major groups that emerged from these studies are the host nucleic acid metabolism proteins that modulate translation and/or replication of the viral RNA, and membrane metabolism proteins that are hijacked to support the structural and functional development of the viral replication organelles, specialized membranous platforms harboring the viral replication complexes. Currently, neither the stoichiometry of the viral proteins nor the full spectrum of the cellular factors required for the activity of the enterovirus replication complexes are known.

Multiple cellular factors can directly interact with specific structural elements of viral RNA and affect its stability, translation, and replication efficiency. Importantly, while the enterovirus life cycle takes place exclusively in the cytoplasm, many of such proteins are normally restricted to the cellular nucleus. Enteroviruses gain access to the nuclear proteins through the action of a protease 2A which specifically cleaves several nucleoporins resulting in the inactivation of organized nucleo-cytoplasmic trafficking and release of the nuclear proteins into the cytoplasm (18-21). A few well-studied examples include a DNA repair component tyrosyl-DNA phosphodiesterase 2 (TDP2) which removes the Vpg from the 5’ end of the viral RNA associated with polysomes (22, 23). Nuclear mRNA processing factors polypyrimidine tract binding protein 1 (PTB1) and poly(rC) binding proteins 1 and 2 (PCBP1 and PCBP2) interact with the IRES elements of different picornaviruses, including enteroviruses, and stimulate translation and/or replication of the viral RNAs (22, 24, 25). Heterogeneous nuclear ribonucleoprotein C (HNRNPC) is required for optimal functioning of the poliovirus RNA replication complex (26). Over the years, several other host nuclear proteins have been identified that interact with specific structural elements of the RNAs of enteroviruses and other picornaviruses, however, the full extent of the effects exerted by the complex mixture of the nuclear RNA-binding factors that are translocated to the cytoplasm upon infection is still far from understood (27, 28). Interestingly, the accumulating evidence demonstrates that none of such RNA binding and/or processing proteins is absolutely required for the infection, suggesting a significant redundancy of the host protein functions in supporting viral RNA translation/replication cycle (22, 23, 29, 30). Rather, the cumulative effect of multiple host proteins on the viral RNA stability, translation, and replication efficiency is likely cell type-dependent, contributing to the determination of the viral tropism in the host (31, 32).

Viral RNA replication and, likely, translation, especially during the later stages of infection, are associated with replication organelles. These structures feature unique membrane and protein composition, and their establishment and expansion depend on the virus-induced reconfiguration of the cellular lipid and membrane synthesis and trafficking pathways. Accordingly, several key host proteins hijacked by enteroviruses have been identified that are responsible for the structural development and the acquisition of a specific replication-conducive biochemical signature of the replication organelles. CTP-phosphocholine-cytidyl transferase alpha (CCTα), the rate-limiting enzyme in the phosphatidylcholine synthesis pathway, lipid droplet-associated lipases adipocyte triglyceride lipase (ATGL) and hormone-sensitive lipase (HSL), as well as several long-chain acyl-CoA synthetases (ACSLCs), are implicated in the activation of infection-specific phospholipid synthesis that provides the bulk of membrane material for the expansion of the replication organelles (33-35). Recruitment of GBF1, a guanine nucleotide exchange factor for small GTPases Arf (ArfGEF), results in a massive association of Arfs with the replication organelles. Phosphatidylinositol 4 kinase III beta (PI4KIIIβ) and oxysterol binding protein (OSBP) together with several other factors mediate the enrichment of the replication organelles in phosphatidylinositol 4 phosphate (PI4P) and cholesterol. Inhibition of GBF1, PI4KIIIβ or OSBP activities severely restricts the replication of diverse enteroviruses (36-50). By the end of the enterovirus replication cycle, the replication organelles may occupy most of the cellular volume (51, 52). Thus, these membranes should be significantly enriched in the factors that support the translation and/or replication of the viral RNA, but also likely in those that may possess a direct anti-viral activity.

Here we took advantage of a strict dependence of enterovirus replication on a cellular protein GBF1 to perform a proteomics characterization of the replication organelles. GBF1 is recruited to the replication organelles through direct interaction with the enterovirus non-structural protein 3A, and the ArfGEF activity of GBF1 is required to support the viral RNA replication (46-48). Thus, GBF1 is likely localized on the replication organelles close to the active replication complexes. GBF1 is a large multi-domain protein normally engaged in multiple protein-protein and protein-membrane interactions (reviewed in (53)). We previously demonstrated that the C-terminal part of GBF1 is dispensable for enterovirus replication (54, 55). To reduce the background of the proteins that are not likely to be important for viral replication, we used such a C-terminally truncated GBF1 to generate a fusion with the APEX2 peroxidase. This peroxidase in the presence of H_2_O_2_ and biotin-phenol generates short-lived active biotin-phenoxyl radicals that covalently attach to the electron-rich amino-acids of nearby proteins. The APEX2-based proximity biotinylation assay has been successfully used for the characterization of proteomes of different compartments of eukaryotic cells (56-58). In non-infected cells, the truncated APEX2-GBF1 construct diffusely localized in the cytoplasm in non-infected cells, but was effectively recruited to the replication organelles and was fully functional in supporting poliovirus replication. Accordingly, the profile of the biotinylated proteins isolated from mock- and poliovirus-infected cells was significantly different. Among the biotinylated viral proteins, i.e. those localized close to GBF1, the intermediate products of polyprotein processing were significantly enriched, suggesting that either the GBF1 environment is associated with active polyprotein processing, or that those incomplete products of proteolysis may perform specific functions in the GBF1-enriched domains of the replication organelles. The largest group of host proteins identified in infected samples were those involved in RNA metabolism, many of which are normally localized in the nucleus, underscoring the massive relocalization of nuclear proteins upon infection and their engagement in the replication process. Many of these proteins have been previously reported to be associated with the replication of diverse enteroviruses, validating our approach. Knock-down of expression of several of the most abundant proteins identified in our assay revealed both pro- and anti-viral factors, affecting translation/replication step of the viral RNA life cycle. Interestingly, one of the strongest negative effects on viral replication was observed upon the knock-down of expression of fructose-bisphosphate aldolase A (AldoA), a glycolytic enzyme important for the ATP biogenesis and the production of ribose-5- phosphate, necessary for *de novo* nucleotide synthesis. We observed similar effects of the depletion of the assayed proteins on the development of infection of poliovirus and Coxsackie virus B3, representatives of the Enterovirus C and B species, respectively, indicating a conservation of the enterovirus replication mechanisms.

Thus, our data significantly expand the repertoire of the known cellular proteins involved in the development of enterovirus infection and elucidate the complex composition of pro- and anti- viral factors associated with the replication organelles.

## Materials and methods

### Cells and viruses

HeLa and RD cells were maintained in DMEM high glucose modification supplemented with pyruvate and 10% FBS. Retrovirus packaging cell line GP2-293 was maintained in DMEM high glucose modification supplemented with 10% FBS. Cell viability was determined using either CellTiter-Glo kit (Promega) or XTT assay (Thermo Fisher) that detect the level of cellular ATP, or the activity of the mitochondrial respiratory chain enzymes, respectively, according to the manufacturers’ recommendations. Poliovirus type I (Mahoney) and Coxsackie virus B3 (Nancy) were rescued using the plasmids pXpA and p53CB3/T7 coding for the full-length viral cDNAs under the control of a T7 promotor kindly provided by Dr. Raul Andino (University of California, San Francisco) and Prof. van Kuppeveld (University of Utrecht, the Netherlands), respectively. The viruses were propagated in HeLa cells, viral iters were determined by plaque or TCID_50_ assays on RD cells grown on 6- or 96-well plates, respectively, using 10x dilutions of the virus preparations. TCID_50_ titers were calculated using Kärber’s formula (59).

### Plasmids

APEX2 coding sequence (56) with the FLAG epitope at the N-terminus optimized for expression in human cells was synthesized by Invitrogen (GeneArt service). For the transient expression, the FLAG-APEX2 was fused in-frame upstream of the GBF1 truncated after HDS1 domain and containing a BFA-resistant Sec7 domain from ARNO (GARG-1060) in a mammalian expression vector pCI (Promega) generating a plasmid pCI-FLAG-APEX2-GARG- 1060. For stable expression, the FLAG-APEX2-GARG-1060 insert was cloned into the retroviral vector plasmid pLNCX2 (Takara Bio). Cloning details are available upon request. Plasmid pEGFP-GARG-1060 coding for the truncated GBF1 with a BFA-resistant Sec7 domain under the control of a CMV promotor was described in (54). Plasmids pXpA-RenR and pRib-CB3- RLUC coding for cDNAs of polio and Coxsackie B3 replicons with Renilla luciferase substituting the capsid coding region P1 under the control of a T7 promotor were described in (60, 61).

### Establishment of a HeLa cell line stably expressing APEX2-GBF1 construct

Retrovirus packaging cell line GP2-293 (Takara Bio) expressing Moloney murine leukemia virus gag and pol genes was co-transfected with pLNCX2 vector with the FLAG-APEX2-GARG1060 insert and a plasmid coding for the vesicular stomatitis virus envelope glycoprotein (Takara Bio) using Mirus 2020 DNA transfection reagent (Mirus). The infectious virions were harvested in the culture supernatant 48 h post-transfection. HeLa cells seeded into a 6-well plate were transduced with the freshly produced retrovirus virions in the complete growth medium supplemented with 10 µg/ml Polybrene (Millipore Sigma). The plate was centrifuged at 1,200g for 1 h at 32°C to enhance the transduction efficiency and kept at 37C for 18 h. The next day, the transduction medium was replaced with a fresh complete growth medium, and cells were incubated overnight. Forty-eight hours after the start of transduction, cells were transferred into a T-25 flask, and the drug-resistant colonies were selected with 300 µg/ml G418 (VWR Life Science) for two weeks. The resistant colonies were pooled, and the stable cell lines were maintained in the complete growth medium supplemented with 300 µg/ml G418. At this point, approximately 60% of the cells showed the expression of the transgene. For a clonal selection, the cells were seeded at a density of ∼0.3 cell/well in a 96 well plate and the colonies established from the individual cells were screened for the transgene expression. A colony that showed >90% uniform pattern of a functional transgene expression as demonstrated by anti-FLAG staining, biotinylation reaction, and polio replicon replication in the presence of 2µg/ml of brefeldin A (Millipore Sigma) was expanded and used for the rest of the study.

### Antibodies

*Cellular proteins:* Mouse monoclonal antibodies: anti-GBF1 (BD Biosciences (612116)), anti-EWSR1 (Novus Biologicals (NBP1-92686)), anti-AldoA (ProteinTech (67453-1- Ig)), anti-ACBD3 (Millipore Sigma (SAB1405255)), anti-β-actin antibodies conjugated with horseradish peroxidase (HRP) (Millipore Sigma (A3854)). Rabbit polyclonal antibodies: anti-OSBP (ProteinTech (11096-1)), anti-PI4KIIIβ (Millipore Sigma (06-578)), anti-ILF3-90 (Millipore Sigma (HPA001897)), anti-FLAG tag (Thermo Fisher (PA1-984B)).

*Viral antigens:* Mouse monoclonal anti-poliovirus VP3, 2B, 2C, and 3A, and rabbit polyclonal antibodies against poliovirus 3B were a gift from prof. Kurt Bienz (University of Bazel, Switzerland) and were partially described in (62-64). Rabbit polyclonal anti-poliovirus 3D antibodies were custom generated by Chemicon and were described in (35). Mouse monoclonal anti-dsRNA antibody J2 was from English & Scientific Consulting Kft.

Secondary Alexa dye fluorescent antibody and streptavidin conjugates were from Thermo Fisher, HRP secondary antibody conjugates were from Amersham or KPL.

### Biotinylation reaction

Depending on the future analysis, HeLa cells stably expressing FLAG- APEX2-GARG1060 were seeded in either a 12-well plate with or without coverslips, a T-25, or a T-225 flask. The cells were infected (or mock-infected) with 10 PFU/cell of poliovirus, and incubated in the presence of 2 μg/ml BFA. At 30 min before the indicated times post-infection, the incubation medium was replaced with pre-warmed DMEM with 500 μM biotinyl tyramide (biotin-phenol) (Chemodex). For the biotinylation reaction, the medium was replaced with PBS containing 20 mM H_2_O_2_ (Millipore Sigma) and incubated for three min at 37°C. Then the cells were washed three times with PBS and either immediately fixed with 4% formaldehyde (Electron Microscopy Sciences) in PBS for microscopy, or lysed in RIPA lysis buffer supplemented with a proteinase inhibitor cocktail (Millipore Sigma) followed by sonication. The sonicated RIPA lysates were used either directly for SDS-PAGE and western blotting, or for further purification of biotinylated proteins.

### Purification of biotinylated proteins

The whole-cell lysates were mixed with the streptavidin magnetic beads (Pierce) equilibrated in RIPA buffer and incubated on a rotator for 1 h at room temperature. The beads were collected using a magnetic rack and washed three times with RIPA buffer. The bound proteins were eluted by boiling the beads in 40 µL of 3X Laemmli sample buffer supplemented with 2 mM biotin and 20 mM dithiothreitol for 10 min. The beads were removed using a magnetic rack, and the eluates were kept at -80°C for further analysis.

### siRNAs

The following sense siRNA sequences targeting the expression of human genes were used:

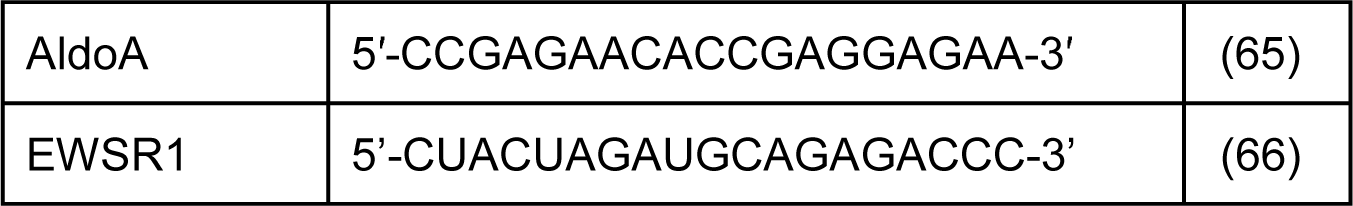

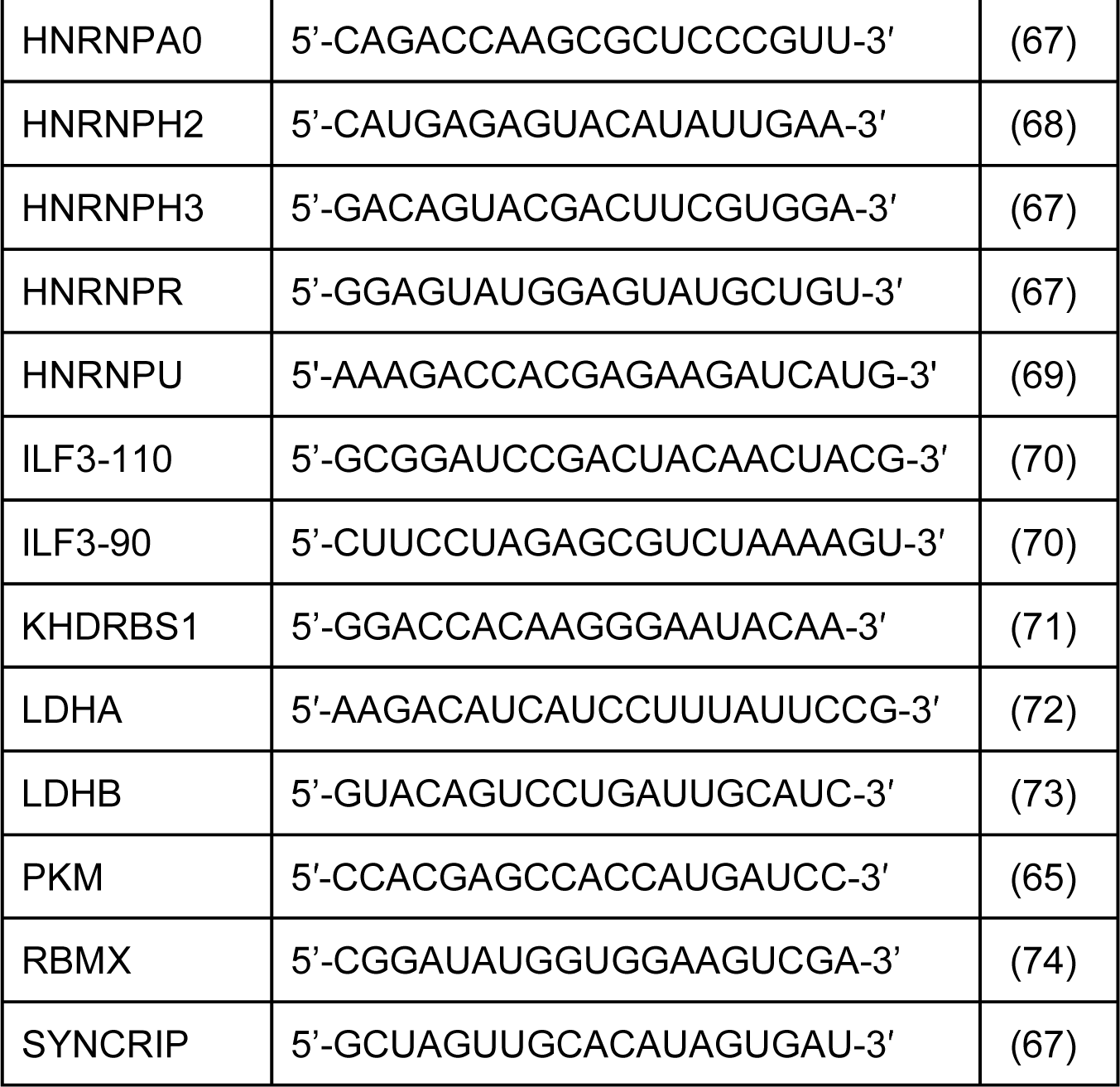

The siRNA duplexes were synthesized with 3’ UU overhangs by Dharmacon. The siRNAs were transfected into HeLa cells using Dharmafect 1 transfection reagent according to the manufacturer’s protocol, and the experiments were performed ∼72h after siRNA transfection. As a non-targeting control siControl#1 (Ambion) was used.

### Microscopy

Cells grown on a coverslip in a 12 well plate were fixed with 4% formaldehyde (Electron Microscopy Sciences) in PBS for 20 min, washed three times with PBS and permeabilized for 5 min in 0.2% Triton-X100 (Millipore Sigma). The cells were incubated sequentially with the primary and secondary antibodies diluted in PBS with 3% blocking reagent (Amersham) for 1 h each with 3x PBS washes in between. Confocal images were taken with Zeiss LSM 510 microscope under the control of ZEN software (Zeiss). All images from one experiment were taken under the same microscope settings. Structural illuminated microscopy super-resolution (SIM) images were taken with Nikon A1R microscope under the control of NIS- Elements software (Nikon). Digital images were processed with Adobe Photoshop software for illustrations, all changes were applied to the whole image, and images from the same experiments were processed with the same settings.

### Replicon assay

Replicon assays were performed essentially as described in (60). Briefly, purified replicon RNA was transfected into HeLa cells grown on 96 well plates using TransIT mRNA transfection reagent (Mirus), and the cells were incubated in the growth medium supplemented with 5µM of cell-permeable Renilla luciferase substrate EnduRen (Promega) at 37C directly in an ID3 multiwell plate reader (Molecular Devices). The measurements were taken every hour for 18 hours, the data were processed using GraphPad Prism software. Total replication was calculated as the area under the curve for the kinetic luciferase measurement for each well, the signal from at least 16 wells was averaged for each sample, unpaired t-test was used to compare the differences within pairs of experimental and control conditions, p<0.05 was considered significant.

### Proteomics analysis

Biotinylated proteins collected from infected (or mock-infected) HeLa cells grown on a 225cm^2^ flask (approximately 3E7 cells/flask) from five independent experiments were pooled together and run into 12% SDS-PAGE for ∼1cm. The gel was stained with Coomassie and the gel slices containing proteins from infected and mock-infected cells were excised and processed for proteomics analysis at the Harvard proteomics facility as follows:

*Sample Preparation procedure:* Gel slices were washed in 50 % acetonitrile and rehydrated with 50 mM ammonia bicarbonate trypsin solution for overnight digestion at 37C. The next day peptides were extracted with a series of elution and completely dried down in a speed vac. The samples were solubilized in 0.1 % formic acid in water for analysis by tandem mass spectrometry.

*Mass spectrometry analysis:* The LC-MS/MS experiment was performed on a Lumos Tribrid Orbitrap Mass Spectrometer (Thermo Fischer) equipped with Ultimate 3000 (Thermo Fisher) nano-HPLC. Peptides were separated onto a 150µm inner diameter microcapillary trapping column packed first with approximately 2cm of C18 Reprosil resin (5µm, 100 Å, Dr. Maisch GmbH, Germany) followed by PharmaFluidics (Gent, Belgium) 50cm analytical column. Separation was achieved by applying a gradient from 5– 27% acetonitrile in 0.1% formic acid over 90 min at 200 nl/min. Electrospray ionization was enabled by applying a voltage of 2 kV using a homemade electrode junction at the end of the microcapillary column and sprayed from metal tips (PepSep, Denmark). The mass spectrometry survey scan was performed in the Orbitrap in the range of 400–1,800 m/z at a resolution of 6×10^4^, followed by the selection of the twenty most intense ions (TOP20) for CID-MS2 fragmentation in the Ion trap using a precursor isolation width window of 2 m/z, AGC setting of 10,000, and a maximum ion accumulation of 100 ms. Singly charged ion species were not subjected to CID fragmentation. The normalized collision energy was set to 35 V and an activation time of 10 ms. Ions in a 10 ppm m/z window around ions selected for MS2 were excluded from further selection for fragmentation for 60s.

*Data Analysis:* Raw data were submitted for analysis in Proteome Discoverer 2.4 (Thermo Scientific) software. Assignment of MS/MS spectra was performed using the Sequest HT algorithm by searching the data against a protein sequence database including all entries from our Uniport_Human2018_SPonly database as well as other known contaminants such as human keratins and common lab contaminants. Quantitative analysis between samples was performed by label-free quantitation (LFQ). Sequest HT searches were performed using a 10 ppm precursor ion tolerance and requiring N-/C termini of each peptide to adhere to Trypsin protease specificity while allowing up to two missed cleavages. Methionine oxidation (+15.99492 Da), as well as deamidation (+ 0.98402 Da) of Asparagine and Glutamine amino acids, were set as variable modifications. Special modification of 1xBiotin-tyramide (+361.14601 Da) on Tyrosine amino acid residues was used as a variable modification. All cysteines were set to a permanent modification with carbamidomethyl (+ 57.02146 Da) due to an alkylation procedure. All MS2 spectra assignment false discovery rate (FDR) of 1% on both protein and peptide levels was achieved by applying the target-decoy database search by Percolator (75).

*Gene ontology analysis:* The sets of proteins identified in the infected and mock-infected samples were analyzed for Gene Ontology (GO) term enrichment using PANTHER classification system web tool (76) against all *Homo sapiens* protein-coding genes using Fisher’s exact test and Bonferroni correction for multiple testing. Only the statistically significant enrichment results with p<0.05 are reported.

## Results

### Establishment and characterization of an APEX2-GBF1 system for proximity biotinylation

The Arf-activating function of GBF1 and other ArfGEFs is mediated by their Sec7 domains. A fungal metabolite brefeldin A (BFA) inhibits the Sec7 function of GBF1 and some other but not all ArfGEFs (77). Previously, we developed a GBF1 construct containing a Sec7 domain from another cellular ArfGEF, ARNO, which is not sensitive to BFA (GARG, from <u>G</u>BF1- <u>AR</u>NO-<u>G</u>BF1) (78). The advantage of such BFA-insensitive constructs is that the endogenous GBF1 can be inactivated by BFA so that it is possible to explore the GBF1-related functions supported exclusively by the exogenously introduced BFA-insensitive GBF1 derivatives. We also previously established that the C-terminal part of GBF1 downstream of the HDS1 domain is dispensable for viral replication (54, 55). We reasoned that for the proteomics studies of replication organelles such “minimal” GBF1 constructs would be advantageous since the interactions with the C-terminal part of GBF1 that are non-essential for viral replication will be eliminated. Thus, we fused APEX2 peroxidase N-terminally to the GARG truncated at the end of the HDS1 domain (APEX2-GARG-1060 construct, Fig 1A). We also introduced a FLAG epitope at the very N-terminus part of the construct. To see if APEX2 fusion is compatible with the functioning of the GARG construct in viral replication, HeLa cells were transfected with a plasmid expressing APEX2-GARG-1060 construct, EGFP-GARG-1060 (positive control), or an empty vector (negative control). The next day the cells were transfected with a polio replicon RNA expressing a *Renilla* luciferase gene instead of the capsid proteins, and the replication was monitored in the presence and in the absence of BFA. The inhibitor blocked the replication in the cells transfected with an empty vector, but cells expressing either APEX2- or EGFP-GARG fusions similarly supported the replication in the presence of the drug (Fig. 1B, transfection panel). It should be noted that the replication signal in the presence of BFA is detected only from the cells expressing the resistant GBF1 constructs, which is limited by the transfection efficacy. Thus, the APEX2-GARG-1060 construct is fully functional in polio replication.

**Figure 1.**
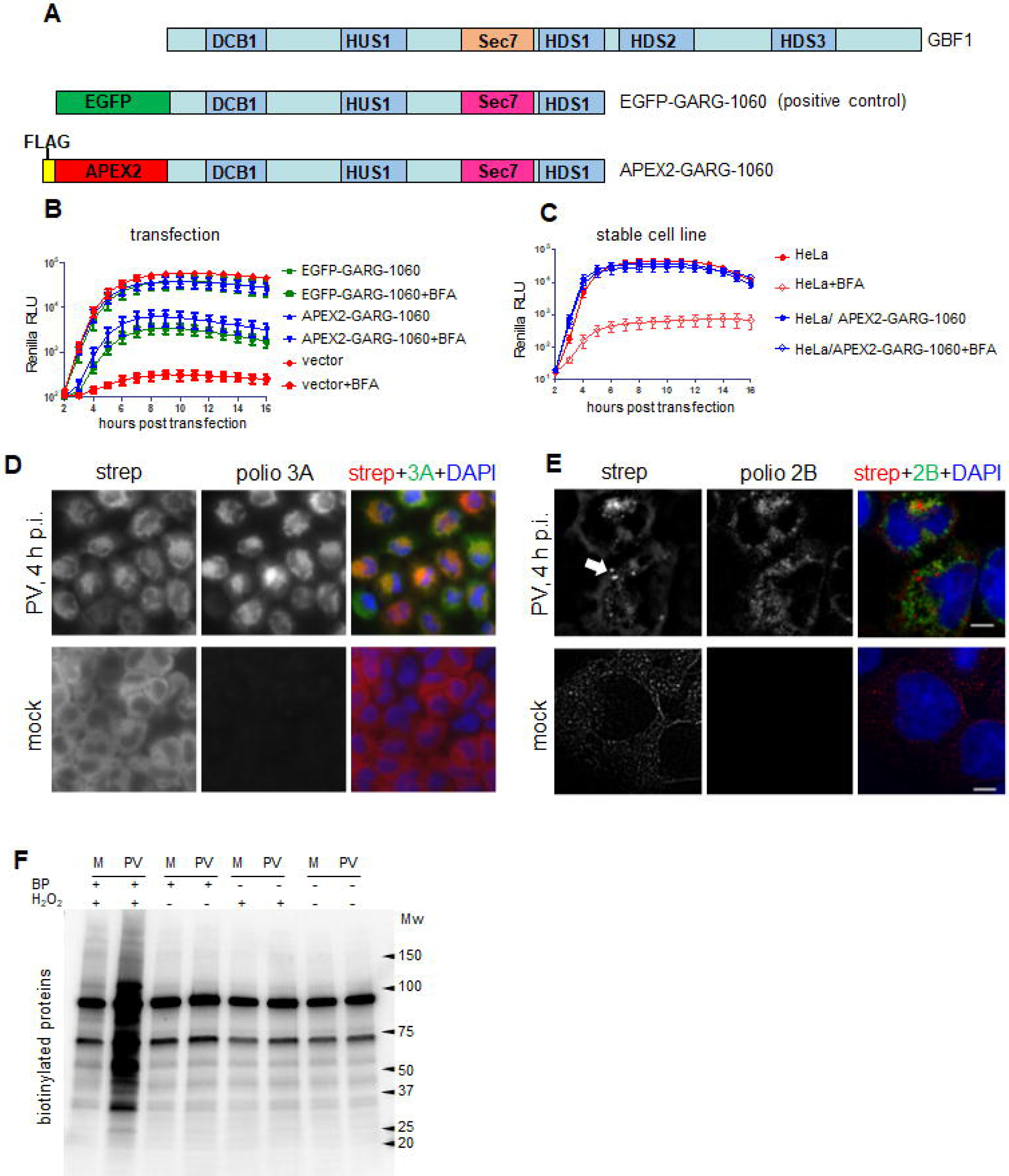
Characterization of the APEX2-GBF1 proximity biotinylation system. **A.** Schemes of the GBF1 domain organization and the C-terminally truncated GBF1 constructs fused to EGFP (positive control) and APEX2. In both GBF1 truncated constructs the cognate Sec7 domain is substituted for a BFA-resistant Sec7 domain from another ArfGEF, ARNO. **B.** HeLa cells were transfected with plasmids expressing the C-terminally truncated GBF1 fusions with APEX2 or EGFP, or an empty vector, and the polio replicon replication was assessed in the presence or absence of 2µg/ml of BFA. **C.** The polio replicon replication assay was performed in the control HeLa cells, or the stable HeLa cell line expressing APEX2-GARG-1060 with or without 2µg/ml of BFA. **D.** The stable HeLa cells line expressing APEX2-GARG-1060 was infected (or mock-infected) with 10 PFU/cell of poliovirus, and the biotinylation reaction was performed at 4 h p.i. The cells were processed for visualization of biotinylated proteins with a fluorescent streptavidin conjugate and staining for a poliovirus antigen 3A. **E.** The stable HeLa cells expressing APEX2-GARG-1060 cells were infected (or mock-infected) with poliovirus and the biotinylation reaction was performed as in D. The cells were stained with a fluorescent streptavidin conjugate and antibodies against a poliovirus antigen 2B and processed for structural illumination superresolution microscopy. The arrow shows round bright biotinylation- positive structures identified as stress granules. The scale bar is 10µm. **F.** The stable HeLa cell line expressing APEX2-GARG-1060 was infected (PV), or mock-infected (M) with 10 PFU/cell of poliovirus, and the specificity of the protein biotinylation was assessed by performing the biotinylation reaction at 4 h p.i. with biotin-phenol (BP) and hydrogen peroxide (complete reaction), or without one, or both of the compounds.

The transiently-transfected cells are not well suited for proteomics studies because the level of transgene expression varies greatly and also because the transfection reagents and presence of a plasmid DNA in the cytoplasm could trigger the innate immune responses. Thus, we established a stable cell line expressing APEX2-GARG-1060 construct by a retrovirus vector transduction and a clonal selection so that the resulting culture expressed a uniform level of the transgene. Polio replicon replicated equally efficiently in these cells in the presence and in the absence of BFA, while in the control HeLa cells BFA blocked polio replicon replication (Fig. 1B, stable cell line panel). This cell line was used for all further experiments which were performed in the presence of BFA so that the replication was supported exclusively by the APEX2-GARG- 1060 construct.

To assess the protein biotinylation, the cells were infected (or mock-infected) with poliovirus at a multiplicity of infection (MOI) of 10 plaque-forming units (PFU)/cell (so that all cells are infected) and the biotinylation reaction was performed at 4 hours post-infection (h p.i.) (in the middle of the poliovirus replication cycle). The cells were stained with a fluorescent streptavidin conjugate to visualize the biotinylated proteins, and with an antibody against a viral non-structural antigen 3A. In mock-infected cells, the streptavidin signal was diffusely distributed in the cytoplasm (although higher magnification images showed some association of the staining with intracellular structures), consistent with the loss of GBF1-specific subcellular targeting of the APEX2-GARG-1060 construct due to the removal of the C-terminal part of GBF1, containing most of the membrane-targeting determinants (79, 80) (Fig. 1D, mock). In infected cells, however, the biotinylation pattern was strictly confined in bright perinuclear rings, the characteristic localization of poliovirus replication organelles, and, accordingly, was extensively co-localized with the 3A signal. It was also visibly brighter than the streptavidin signal in mock-infected cells (Fig. 1D, PV). We also investigated the fine distribution of biotinylated proteins using structural illuminated superresolutiuon microscopy (SIM). The SIM images showed a reticulate pattern in mock-infected cells which is likely mostly due to the mitochondria which are enriched in biotin-containing enzymes (81). In infected cells, a differently structured biotinylation signal was associated with the replication organelles, as evidenced by the staining for the viral antigen 2B (Fig. 1E). Interestingly, we observed in many infected cells bright round foci of biotinylation signal (Fig. 1D, arrow) which were identified as stress granules (see the last section in the Results).

To characterize the specificity of the biotinylation reaction, the cells were similarly infected (or mock-infected), and at 4 h p.i. they were incubated either with both H_2_O_2_ and biotin-phenol, or with H_2_O_2_ or biotin-phenol only, or without any of those compounds. Western blot analysis with streptavidin conjugated to horseradish peroxidase (HRP) showed two major bands of similar intensity in all the samples, likely corresponding to the pyruvate carboxylase and mitochondrial 3-methylcrotonyl carboxylase, biotin-containing enzymes previously observed in studies with APEX2 biotinylation (81). At the same time, only samples incubated in the presence of both compounds showed extensive biotinylation of additional proteins, confirming the specificity of the biotinylation reaction. The level of protein biotinylation in infected cells significantly exceeded that in the mock-infected sample, in accordance with the pattern observed with the staining of cells with a fluorescent streptavidin conjugate.

Thus, the APEX2-GARG-1060 efficiently supports poliovirus replication, it is recruited to the replication organelles and can specifically biotinylate proteins associated with these structures.

### Initial characterization of the biotinylated proteins during the time course of infection. Cellular proteins

An important advantage of APEX2-based biotinylation is a short time of the actual labeling reaction permitting time-resolved snapshots of protein composition. The replication cycle of poliovirus in HeLa cells takes about 6-8 h. We infected cells expressing APEX2-GARG-1060 with 10 PFU/cell of poliovirus and performed the biotinylation reactions at 2, 4 and 6 h p.i. The biotinylated proteins were isolated by streptavidin colomns and analyzed in a western blot assay with a streptavidin-HRP conjugate for the global biotinylation pattern. At 2 h p.i., the amount of the biotinylated proteins and their pattern were similar in infected and control samples, and the mock-infected samples did not significantly change during the time course of the experiment. At 4 and 6 h p.i., protein biotinylation strongly increased in infected cells, and they were distributed through the whole range of molecular weights. Visually, the pattern of the biotinylated proteins from the 6 h p.i. sample was the same as at 4 h p.i., but the amount of the biotinylated proteins was higher (Fig. 2A).

**Figure 2.**
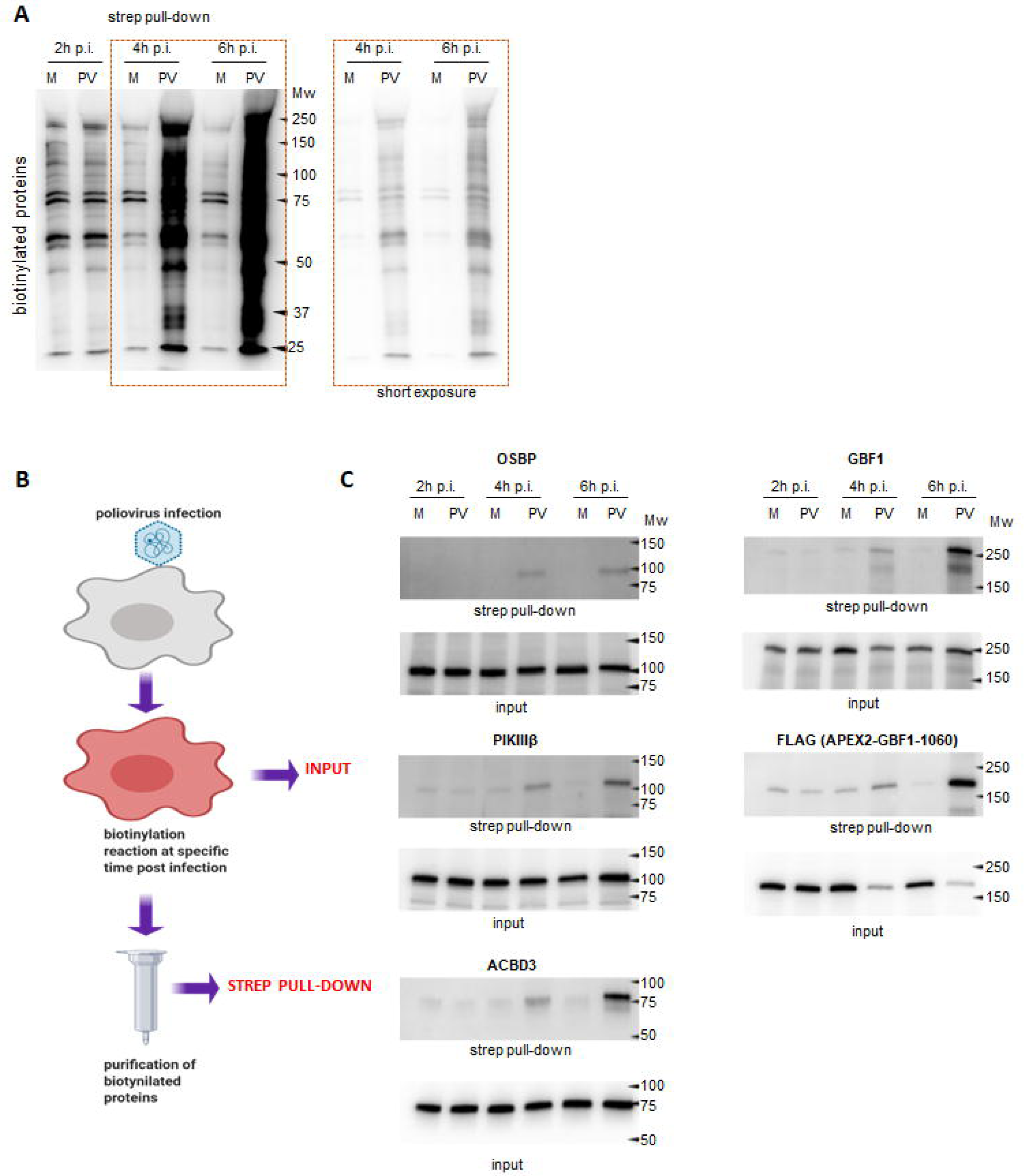
Known cellular proteins recruited to the replication organelles are biotinylated by FLAG-APEX2-GARG1060. **A.** The stable HeLa cell line expressing FLAG-APEX2-GARG- 1060 was infected (PV), or mock-infected (M) with 10 PFU/cell of poliovirus, and the biotinylation reactions were performed at the indicated times post-infection. The biotinylated proteins were collected on streptavidin beads, resolved on SDS-PAGE and analyzed in a Western blot with HRP-conjugated streptavidin. **B.** Scheme of the biotinylation experiment for comparison of the biotinylated protein fraction (strep pull-down) with the total proteins in the cellular lysates (input). **C.** The stable HeLa cells line expressing APEX2-GARG-1060 was infected (PV), or mock-infected (M) with 10 PFU/cell of poliovirus, the biotinylation reactions were performed at 2, 4, and 6 h p.i., and the unfractionated cellular lysates (input) and the biotinylated proteins isolated by streptavidin beads were analyzed with the indicated antibodies against known cellular factors recruited to the replication organelles in a western blot. Anti-FLAG antibodies recognize the APEX2-GARG-1060 protein.

We then analyzed if cellular factors ACDB3, OSBP, and PI4KIIIβ which are involved in the PI4P and cholesterol enrichment of the replication organelles (36, 43, 44, 50) are present in the biotinylated fraction, which would indicate that they localize close to GBF1 in infected cells. The infection and biotinylation reaction were performed as for Fig. 2A. The unfractionated lysates and the proteins recovered in the biotinylated fraction were analysed in western blot with the indicated antibodies (Fig. 2B). We observed a specific increase of the signals for ACDB3, OSBP, and PI4KIIIβ in biotinyulated fraction only in the infected samples collected at 4, and especially 6 p.i. Biotinylated OSBP signal was always observed only in infected samples, while traces of PI4IIIKβ and ACBD3 were also visible in the material recovered from mock-infected samples (Fig. 2C). We also analyzed the biotinylation of endogenous GBF1, and the APEX2- GARG-1060 construct itself. Since the APEX2-GARG-1060 construct lacks the C-terminal part containing the epitope recognized by the anti-GBF1 antibody, it was detected by anti-FLAG antibodies. Again, the strongest signals for both biotinylated GBF1 and APEX2-GARG-1060 were observed in the 4 and 6 h infected samples (Fig. 2C). The increase of the signal for the APEX2-GARG-1060 constructs in 4 and 6 h p.i. samples coincided with a noticeable decrease of the corresponding signal in the total input material, likely reflecting the degradation of the cytoplasmic, but not replication-organelle associated pools of the protein (Fig. 2C). Surprisingly, we did not detect biotinylated Arfs, even though they are enriched on the replication organelles (82, 83), and at least a fraction of Arf molecules is expected to be localized close to GBF1 (data not shown).

### Viral proteins

The proximity biotinylation approach allowed us to identify specific fragments of the poliovirus polyprotein localized in the vicinity of GBF1 on the replication organelles. Since the poliovirus genome is expressed as a single polyprotein undergoing a proteolytic processing cascade, antibodies against a certain antigen will recognize the final maturation product and all the intermediate cleavage products containing this antigen. The available panel of antibodies suitable for western blot (VP3, 2B, 2C, 3A, 3D) covers all known intermediate fragments of the poliovirus polyprotein processing and all individual proteins except capsid proteins VP0 and VP1, proteases 2A and 3C, and the RNA replication protein primer 3B (Vpg) (Fig. 3, poliovirus genome and polyprotein processing scheme). All the tested viral antigens were present in the biotinylated fraction. Interestingly, while in the input material the strongest signals for the viral antigens were found in the final polyprotein cleavage products, in the biotinylated protein fraction the intermediate polyprotein cleavage products were overrepresnted. For example, an uncleaved precursor P2P3 was clearly detected in the biotinylated fraction with anti-3A antibody in 6 h p.i. sample, while this piece of the polyprotein was not visible in the input material (Fig. 3, anti-3A panel). We also observed a specific increase in the biotinylated fraction of the viral antigen-positive fragments that do not correspond to the canonical products of the viral polyprotein processing. Note the red asterisks marking a 3A-positive fragment in the 15-20KDa range (Fig 3, anti-3A panel, 6 h p.i), or a 3D-positive fragment of a molecular weight slightly higher than 3D (Fig. 3, anti-3D panel, 6 h p.i.). This may suggest that the GBF1 environment is associated with active polyprotein maturation, although a preferential enrichment of larger polyprotein fragments due to a higher degree of biotinylation cannot be excluded.

**Figure 3.**
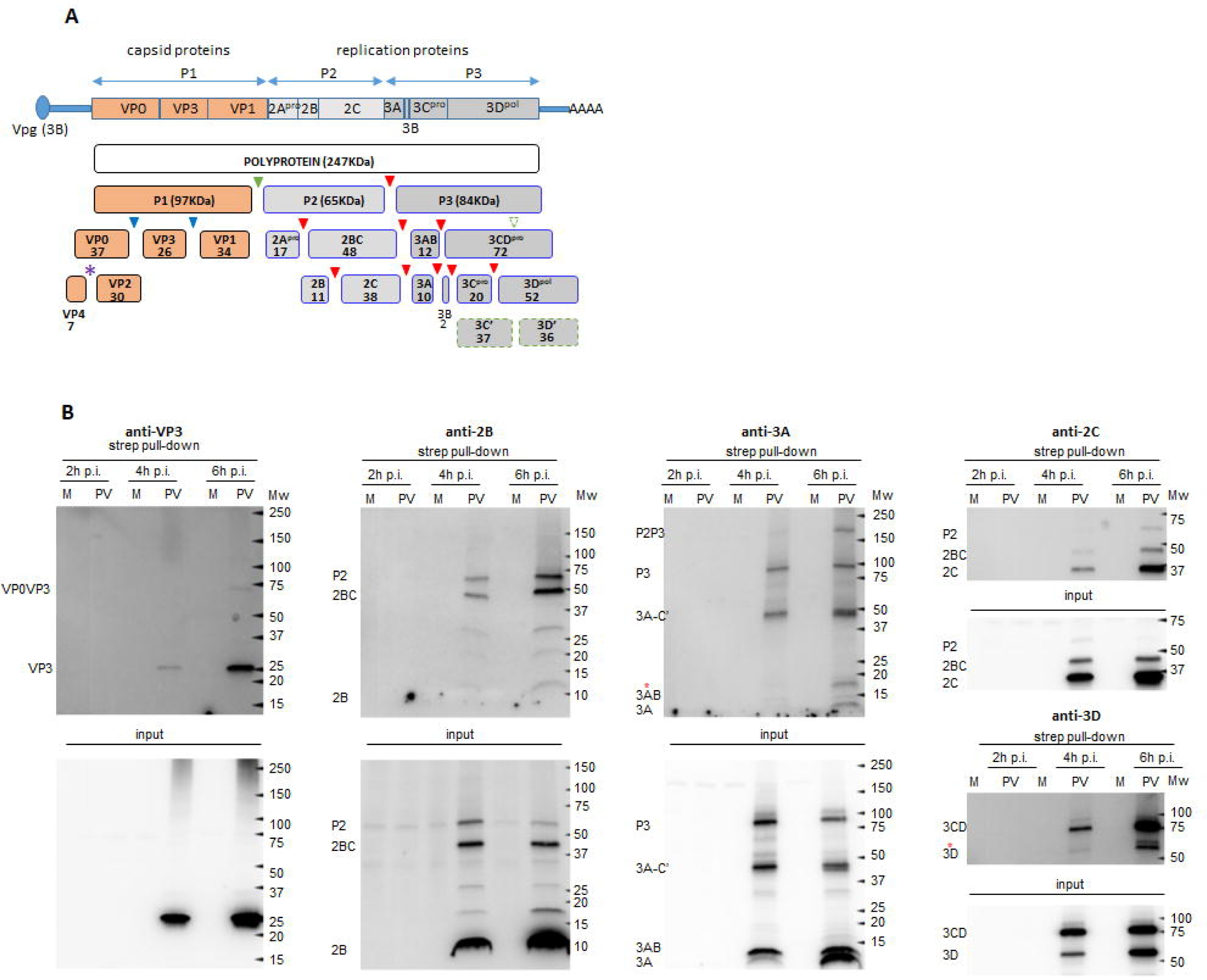
Biotinylation of the viral proteins by APEX2-GARG-1060. **A.** Poliovirus genome and polyprotein processing scheme. The cleavage sites for the viral proteases 2A, 3C, and 2CD are indicated by green, red, and blue filled triangles, respectively. The dashed empty green triangle indicates a 2A cleavage site in 3D believed to be dispensable for replication, the purple star indicates an autocatalytic cleavage site in VP0. **B.** The stable HeLa cell line expressing APEX2-GARG-1060 was infected (PV), or mock-infected (M) with 10 PFU/cell of poliovirus, and the biotinylation reactions were performed at the indicated times post-infection. The biotinylated proteins were collected on streptavidin beads, resolved on SDS-PAGE and analyzed in a Western blot with antibodies against the indicated viral antigens. The antibodies recognize the final and intermediate polyprotein cleavage products containing the corresponding antigen. Red stars on anti-3A and anti-3D panels indicate polyprotein fragments that do not match the known stable polyprotein cleavage products.

Collectively, these results demonstrate that APEX2-GARG-1060 in infected cells can specifically biotinylate both viral and host proteins associated with the replication organelles and that 6h p.i. samples are the most representative for the characterization of the proteome of the replication organelles.

### Proteomics characterization of the replication organelles

For the proteomics analysis, HeLa cells grown on 225 cm^2^ flasks were infected (or mock-infected) with poliovirus at an MOI of 10, the biotinylation reaction was performed at 6 p.i. for 3 min, and the biotinylated proteins were purified on streptavidin colomns. Five independent experiments were performed, and aliquotes of the isolated proteins were assessed in a western blot assay with a streptavidin-HRP conjugate. In all experiments, a similar pattern of a highly increased amount of biotinylated proteins in infected samples was observed, as expected (Fig. 4A). The rest of the purified biotinylated proteins were pooled together and processed for mass-spectrometry protein identification and label-free quantitation (LFQ). Upon filtering the identified proteins from common contaminants, as well as carboxylases which contain naturally covalently attached biotin, and proteins with peroxidase activity which can likely be biotinylated independently of APEX2, 369 and 43 proteins were enriched in the infected and non-infected sets, respectively. 192 proteins in the infected sample and 37 proteins in the mock-infected sample were detected from 2 or more peptides. 331 proteins were found exclusively in the infected sample, while just 5 proteins from the mock-infected sample were not identified in the infected sample (Supplementary Data 1). Among the cellular proteins we previously confirmed to be present among the biotinylated pool by the western blot analysis (see Fig. 2), GBF1 (Q92538) was identified from a total of nine peptides, five of them unique (9 total:5 unique) (further on this designation is used for peptides detected for each protein), while ACBD3 (Q9H3P7) and OSBP1 (P22059) proteins were identified from one peptide each, and PI4KIIIβ (Q9UBF8) was not found (Supplementary Data 1).

**Figure 4.**
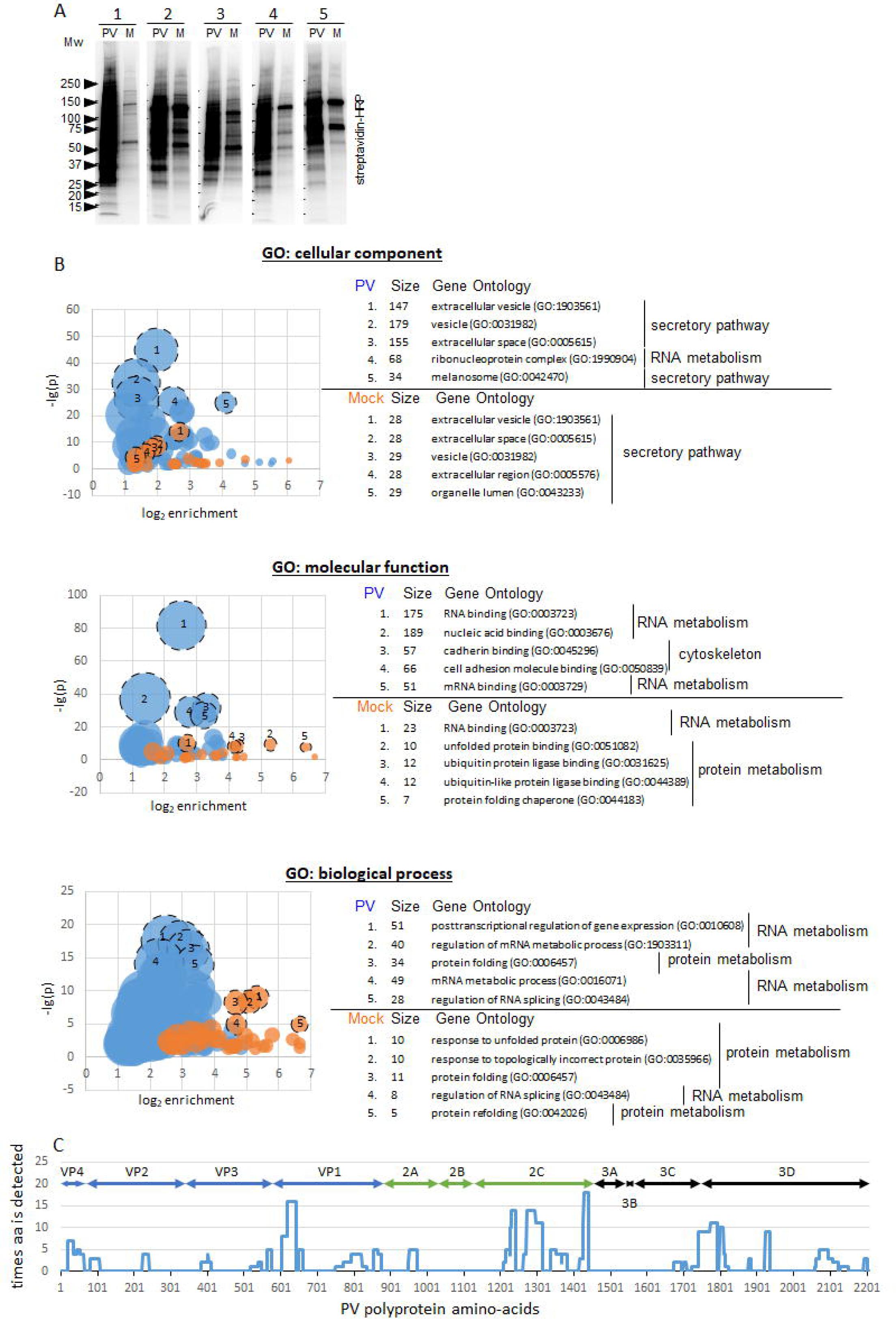
Proteomics characterization of the proteins biotinylated by FLAG-APEX2- GARG1060. **A.** In five independent experiments, the stable HeLa cell line expressing APEX2- GARG-1060 was infected (PV), or mock-infected (M) with 10 PFU/cell of poliovirus, and the biotinylation reaction was performed at 6 h p.i. The biotinylated proteins were purified by streptavidin beads and analyzed in a Western blot with HRP-streptavidin. The biotinylated proteins from these five infected and mock-infected samples were pooled for further proteomics analysis. Full proteomics data is available in Supplementary data 1. **B.** Gene ontology (GO) enrichment analysis of the proteomics data using PANTHER classification system (76). Buble graphs show the number of proteins associated with a particular GO term (bubble size), the log_2_ of enrichment over the expected non-specific associations of genes in the dataset with a particular GO term (x-axis), and the statistical significance of the observed enrichment (negative log_10_ of p-value, y-axis). The five of the most statistically significantly enriched GO terms for proteins from infected and mock-infected samples are shown. The full GO analysis is available in Supplementary data 2. **C.** The distribution of the poliovirus-specific peptides identified by mass-spectrometry analysis over the poliovirus polyprotein. The x-axis shows amino-acid positions in the poliovirus polyprotein, the y-axis shows how many times a particular amino-acid is detected.

Since the proteins were purified upon biotinylation by a GBF1-based construct, one would expect the presence of at least some known interactors of GBF1 or GTPses Arf. Indeed, analysis of the proteins using the Biogrid database of curated interaction data (84, 85) identified 17 Arf1, one Arf3, four Arf4, three Arf5, and four Arf6 interactors. Arf3, Arf4 and Arf5 as well as 12 Arf1 and two Arf6 interactors were identified exclusively among the proteins from the infected sample. Also, 44 proteins were identified as GBF1 interactors, 34 of which were found only in the infected sample (Supplementary Table 1).

The global association of the proteins with cellular structures and pathways was analyzed by Gene Ontology (GO) term enrichment in the cellular component, molecular function, and the biological process categories using the PANTHER classification system (76). In general, the GO term enrichment of the proteins from the infected sample demonstrated a much higher statistical significance than those from the mock-infected cells, which is to be expected given the difference in the number of proteins in each sample. In both samples, the most statistically significantly enriched categories included proteins associated with the cellular secretory pathway and the chaperon-assisted protein folding. In the mock-infected sample, proteins associated with the proteasome-dependent protein degradation were among the highly enriched. In the infected sample, a significant amount of proteins were also associated with the cytoskeleton function. Yet, the nucleic acid, and in particular RNA metabolism, emerged as the predominantly enriched GO terms from the infected sample (Fig. 4B and Supplementary Figure 2). Interestingly, 17 proteins were associated with dsRNA binding, 10 of which were present only in the infected sample (Supplementary Table 1).

The literature search revealed that 62 of the proteins identified in the infected sample were previously reported to have a functional significance for the replication of different enteroviruses, while 50 more were detected in high-throughput screens as proteins that undergo some changes upon enterovirus infection, or as interacting partners with the viral proteins, but their functional significance was not investigated (Supplementary Data 3). The known constituents of the poliovirus replication complex, KH domain-containing, RNA-binding, signal transduction-associated protein 1 (KHDRBS1, Sam68) (86) was identified from 8 total: 2 unique; splicing factor, proline- and glutamine-rich (SFPQ) (27) from 10 total: 6 unique, and polyadenylate-binding protein 1 (PABCP1) (87) from 3 total: 2 unique peptides, exclusively in the infected sample. 6 total:3 unique peptides shared between poly-(rC)-binding proteins 1 and 2 (PCBP1, 2) (88-90) were identified in the infected sample while one unique peptide for each PCBP1 and PCBP2 was detected in the mock-infected control. Polypyrimidine tract-binding protein 1 (PTB1) involved in the activation of the enterovirus IRES-driven translation (91, 92) was detected by one peptide in both infected and mock-infected samples (Supplementary Data 1).

The poliovirus-specific peptides (207 total: 64 unique) identified by the mass-spectrometry analysis were distributed along the whole viral polyprotein, with an intriguing absence of peptides covering 2B and 3A-3B regions. An increased clustering of the detected peptides was observed in the N-terminus of a capsid protein VP1, 3C-3D junction, and in particular in the 2C region (Fig. 4C). This pattern is in accordance with the data shown in Fig. 3 and confirms that the GBF1 environment on the replication organelles is enriched in all viral structural and replication proteins.

Overall, these data validate the relevance of the identified cellular proteins for poliovirus replication and provide an important insight into the viral and cellular protein environment of the replication organelles in the surroundings of GBF1.

### Identification of novel factors affecting viral replication

One of the major goals of this study was to identify novel host factors important for viral replication. The selection of the proteins from a large proteomics dataset for analysis is inevitably arbitrary, but our general criteria were that the proteins should be detected from multiple peptides in the infected sample (i.e. highly enriched), with the previously uncharacterized role in enterovirus replication, and representing different functional clusters.

We selected the following groups: 1. Glycolytic enzymes. Fructose-bisphosphate aldolase A (AldoA), pyruvate kinase (PKM), L-lactate dehydrogenase chain A and B (LDHA and LDHB)) detected from (32 total:15 unique), (15 total:11 unique), (6 total:2 unique) and (4 total: 2 unique) peptides in the infected sample, respectively. Apart from being highly enriched, the group of glycolytic enzymes was selected because LDHA and LDHB are reported Arf interactors and because the glycolytic pathway provides substrates for *de novo* nucleotide synthesis, which may be important for rapidly replicating RNA viruses (84, 85, 93-95).

2. The highly enriched RNA binding proteins. Heterogeneous nuclear ribonucleoproteins A0, H2, H3, R, U (HNRNPA0, H2, H3, R, U), heterogeneous nuclear ribonucleoprotein Q (SYNCRIP), Ewing Sarcoma Breakpoint Region 1 (EWSR1), and RNA-binding motif protein, X chromosome (RBMX) were among the most enriched in the infected sample (10:3, 15:6, 8:5, 10:6, 27:11, 8:6, 12:5, and 7:6 of total: unique peptides, respectively). SYNCRIP and HNRNPU were previously reported in a proteomics screen detecting proteins bound to poliovirus RNA. The depletion of HNRNPU did not affect the virus yield, the contribution of SYNCRIP was not analyzed (96).

3. A potential antiviral factor. A dsRNA binding protein ILF3 was identified from (8 total:4 unique) peptides in the infected sample. This protein was shown to be important for the establishment of dsRNA-induced anti-viral signaling and to either promote or inhibit the replication of diverse viruses (97-101). The ILF3 gene is expressed as multiple isoforms of the two major variants of 90KDa and 110KDa proteins. Both 90KDa and 110KDa proteins have two dsRNA binding domains, with an extended C-terminal GQSY-reach region in the latter (102). A 90KDa isoform of ILF3 was demonstrated to inhibit translation of a poliovirus-rhinovirus chimera RNA in a cell-type dependent manner by binding to the rhino- but not poliovirus IRES (103).

We screened the effects of the siRNA-mediated depletion of these proteins on polio replicon replication and the accumulation of the viral protein 2C upon infection. In the replicon assay, the RNA is transfected into the cells bypassing the normal virion-receptor mediated delivery, so it reflects the RNA translation-replication step of the viral life cycle, while the accumulation of 2C upon infection also integrates the effects of virion-receptor interaction, penetration and uncoating. As a control for the screening methods, we also included siRNA against KHDRBS1 (Sam68) (identified from 8 total: 2 unique peptides in our proteomics dataset), which was previously shown to bind poliovirus polymerase 3D and promote viral replication (86).

Depletion of Sam68 inhibited polio replication in both replication and infection assays, as expected, thus validating our approach (Supplementary Figure 1). Among all the proteins tested, only depletion of LDHA was toxic to cells, so its specific effect on polio replication was impossible to evaluate in this system. Depletion of PKM, LDHB, SYNCRYP and HNRNPU affected the replication in one of the assays only, suggesting a moderate contribution of these proteins in the virus life cycle, at least in the cell culture conditions. Curiously, depletion of RBMX strongly increased the replicon replication but decreased the accumulation of 2C upon infection. Depletion of AldoA, HNRPA0 and EWSR1 showed consistent negative effects on polio replication in both assays, while that of HNRNPR, HNRNPH2, HNRNPH3, and in particular the 90KDa isoform of ILF3 strongly increased the replication in both assays (Supplementary Figure 1).

Thus, our dataset reveals novel host factors with stimulatory and inhibitory effects on poliovirus replication.

### AldoA, EWSR1 and ILF3-90 similarly control the replication of diverse enteroviruses

The proteins AldoA, EWSR1, and the 90K isoform of ILF3 whose depletion consistently significantly affected polio replication were selected for a more detailed analysis. The siRNA knockdown of expression of AldoA and EWSR1 signficantly inhibited the replication of both polio and Coxsackie B3 replicons, while the stimulatory effect of ILF3-90 depletion was detected only with polio replicon (Fig. 5A, B). In conditions of *bona fide* virus infection, the depletion of AldoA and EWSR1 inhibited, while the depletion of ILF3-90 stimulated the infectious virion yield of both viruses (Fig. 5C). Interestingly, while cells treated with the siRNAs against these proteins did not show any obvious signs of cytotoxicity, the viability assay based on the measurement of ATP showed significantly lower values for AldoA depletion (Fig. 5A, viability). We compared the ATP measurement viability test with that based on the activity of a mitochondrial respiratory chain. The latter test did not detect any difference in the cell viability upon depletion of any protein (Fig. 5D). This suggests that the negative effect of the AldoA depletion on the enterovirus replication may be explained by its requirement for ATP production in HeLa cells.

**Figure 5.**
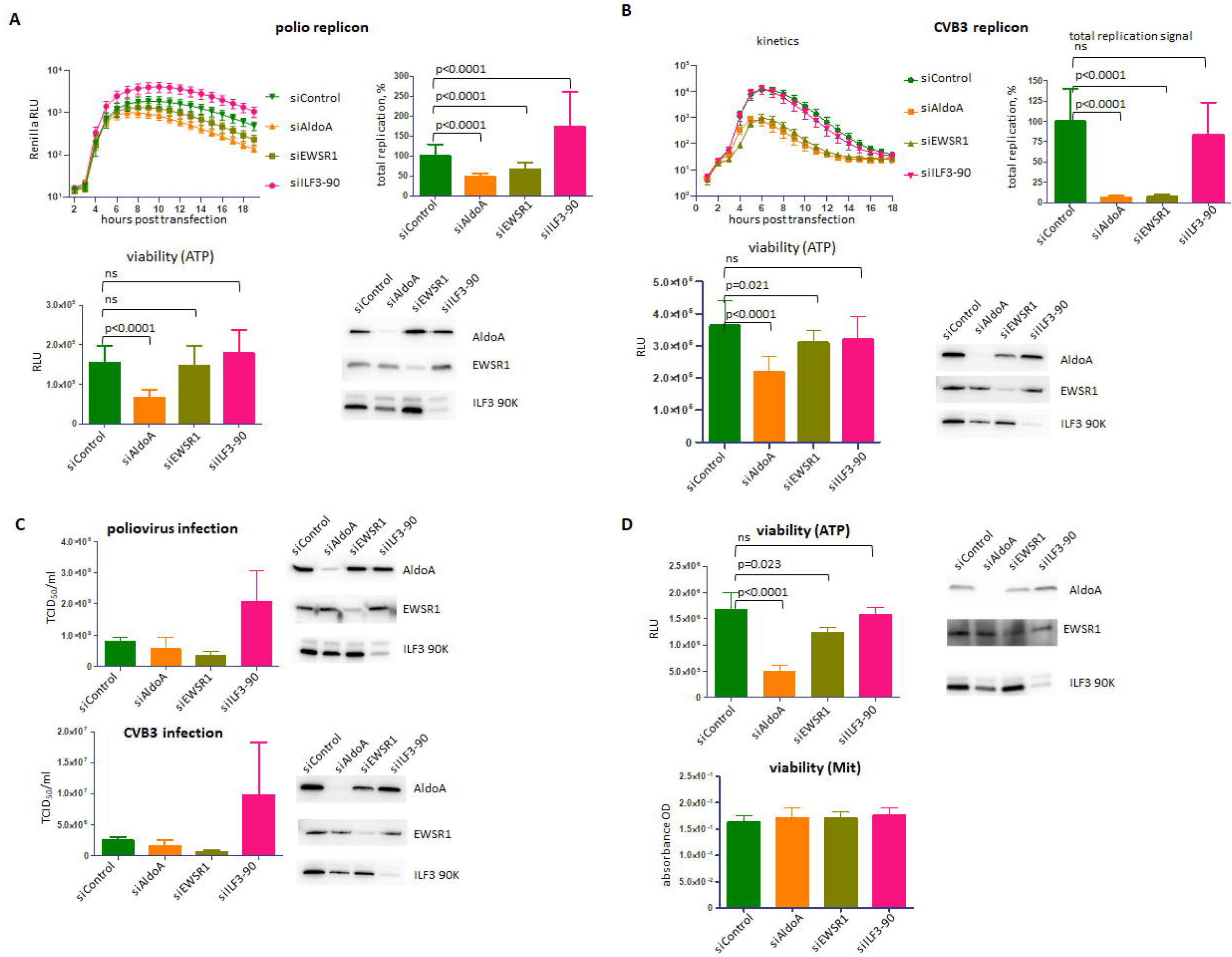
Analysis of the effect of siRNA-mediated knockdown of expression of AldoA, EWSR1 and ILF3-90 on enterovirus replication. **A, B.** HeLa cells were transfected with siRNAs specific against AldoA, EWSR1 and 90KDa isoform of ILF3, or non-targeting control siRNA, and polio or Coxsakie B3 replicon replication assays were performed 72 h post siRNA transfection. The total replication signal was calculated as the area under the corresponding kinetics curves. Cell viability signal is proportional to the level of ATP in cells. Western blots show the efficacy of siRNA-mediated knockdown of the targeted proteins. **C.** HeLa cells were transfected with siRNAs specific against AldoA, EWSR1 and 90KDa isoform of ILF3, or non- targeting control siRNA. 72 h post siRNA transfection the cells were infected with an MOI of 1 PFU/cell of poliovirus or Coxsackie virus B3, and the total virus yield was determined at 6 h p. i. Western blots show the efficacy of siRNA-mediated knockdown of the targeted proteins. **D.** HeLa cells were transfected with siRNAs specific against ALdoA, EWSR1 and 90KDa isoform of ILF3, or non-targeting control siRNA, and the cell viability assays detecting the level of ATP or the activity of the mitochondrial respiratory chain enzymes were performed 72h post siRNA transfection. Western blots show the efficacy of siRNA-mediated knockdown of the targeted proteins.

Finally, we analyzed the cellular distribution of AldoA, EWSR1 and ILF3-90 upon infection. AldoA in mock-infected cells was localized in a dot-like pattern in the cytoplasm, especially in the perinuclear region, and a significant amount of signal was detected in the nucleus, in accordance with the previous report of nuclear accumulation of AldoA in actively dividing cells (104). In poliovirus-infected cells the cytoplasmic AldoA signal was confined within the area of the replication organelles, as evidenced with the staining for the viral membrane-targeted protein 3A (Fig. 6A). Interestingly, Aldo-A positive dots were very closely associated with the signal for dsRNA, but the signals were adjacent, not colocalizing (Fig. 6B).

**Figure 6.**
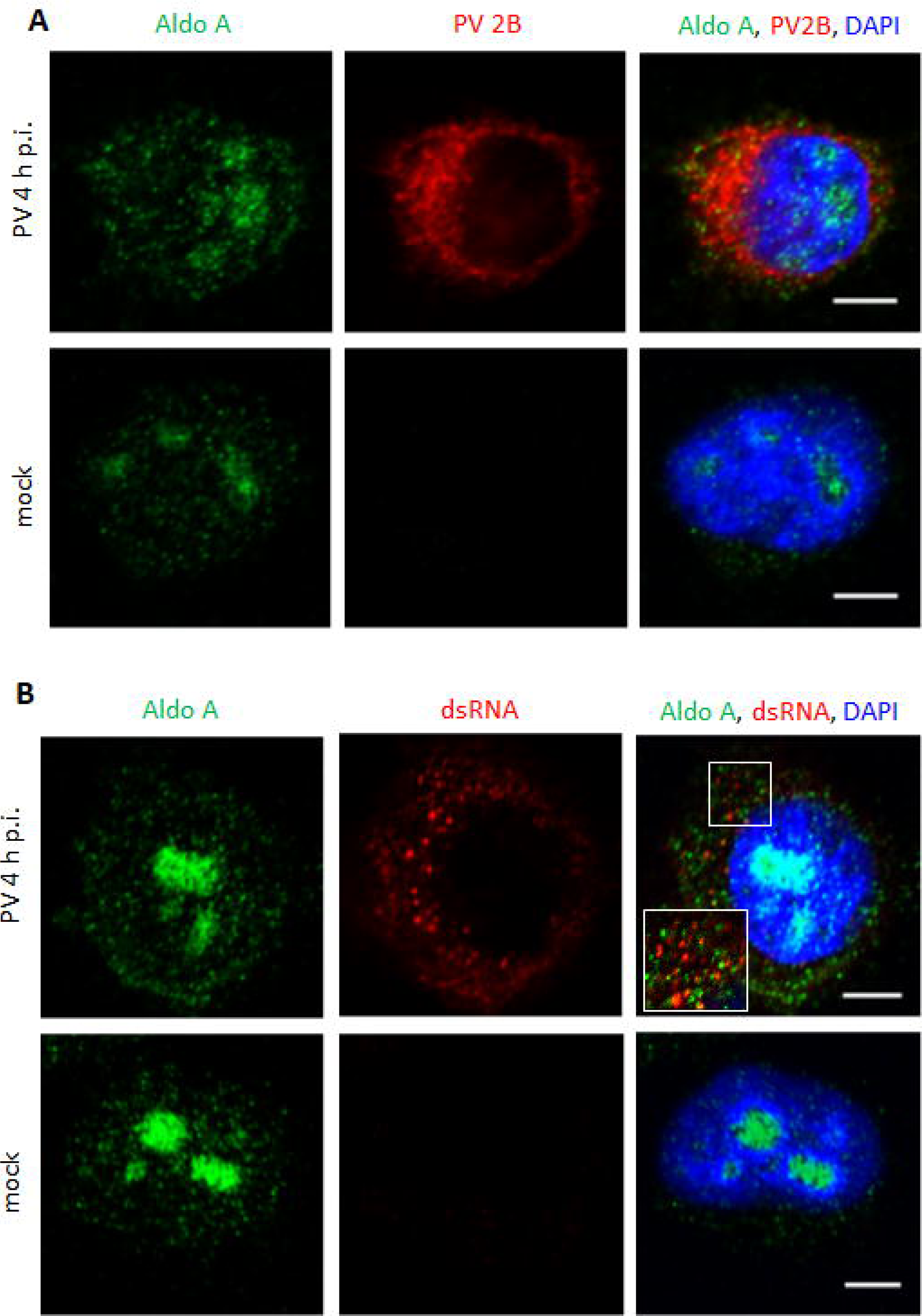
AldoA localization in infected and mock-infected cells. **A, B.** Confocal images of HeLa cells infected (or mock-infected) with 10 PFU/cell of poliovirus, fixed at 4 h p.i. and processed for staining with antibodies against AldoA and a viral antigen 2B, or AldoA and dsRNA, respectively. Scale bar is 10 µm.

EWSR1 in mock-infected cells was found exclusively in the nuclei. In poliovirus-infected cells as early as 2 h p.i. large EWSR1-positive punctae appered in the cytoplasm, and by 4 hp.i., in the middle of poliovirus infectious cycle, all infected cells had exclusively cytoplasmic EWSR1 signal with multiple punctae. By 6 h.p. ESWR1 signal concentrated in the perinuclear area colocalizing with the replication organelles, and the number of punctae per cell was decreased, although they were still clearly detectable in the majority of the cells (Fig. 7A). The cytoplasmic EWSR1 signal in infected cells outside of the punctae strongly colocalized with the structures positive for a viral antigen 3B (Vpg). 3B signal may correspond to the RNA replication primer in a free form, or attached to the 5’ of viral RNAs, but may also be detected as a part of intermediate polyprotein processing products (Fig. 7B). The cytoplasmic punctae pattern of ESWR1 was highly reminiscent of the development of stress granules upon poliovirus infection. To test that, we stained mock-infected and infected cells for EWSR1 and GTPase Activating Protein (SH3 Domain) Binding Protein 1 (G3BP1), a stress granule assembly factor known to be recruited to poliovirus-induced stress granules (105). Indeed, in infected cells, the cytoplasmic punctae of EWSR1 and GRBP1 signals colocolaized perfectly confirming that these structures are stress granules (Fig. 7C). The detection of a stress granule protein upon proximity biotinylation by a GBF1-derived construct was somewhat unexpected since we are not aware of reports of GBF1 targeting to stress granules in infected or otherwise stressed cells. We analyzed the biotinylation pattern relative to the G3BP1 signal in APEX2-GARG-1060 cells. In infected cells we observed multiple bright biotinylation-positive punctae colocalizing with G3BP1-containing stress granules in the cytoplasm, explaining the stress granule protein labeling by APEX2-GARG-1060 construct (Fig. 7D). Whether endogenous GBF1 can be associated with stress granules upon infection, or this is a phenomenon specific to the artificial truncated GBF1 construct requires further investigation.

**Figure 7.**
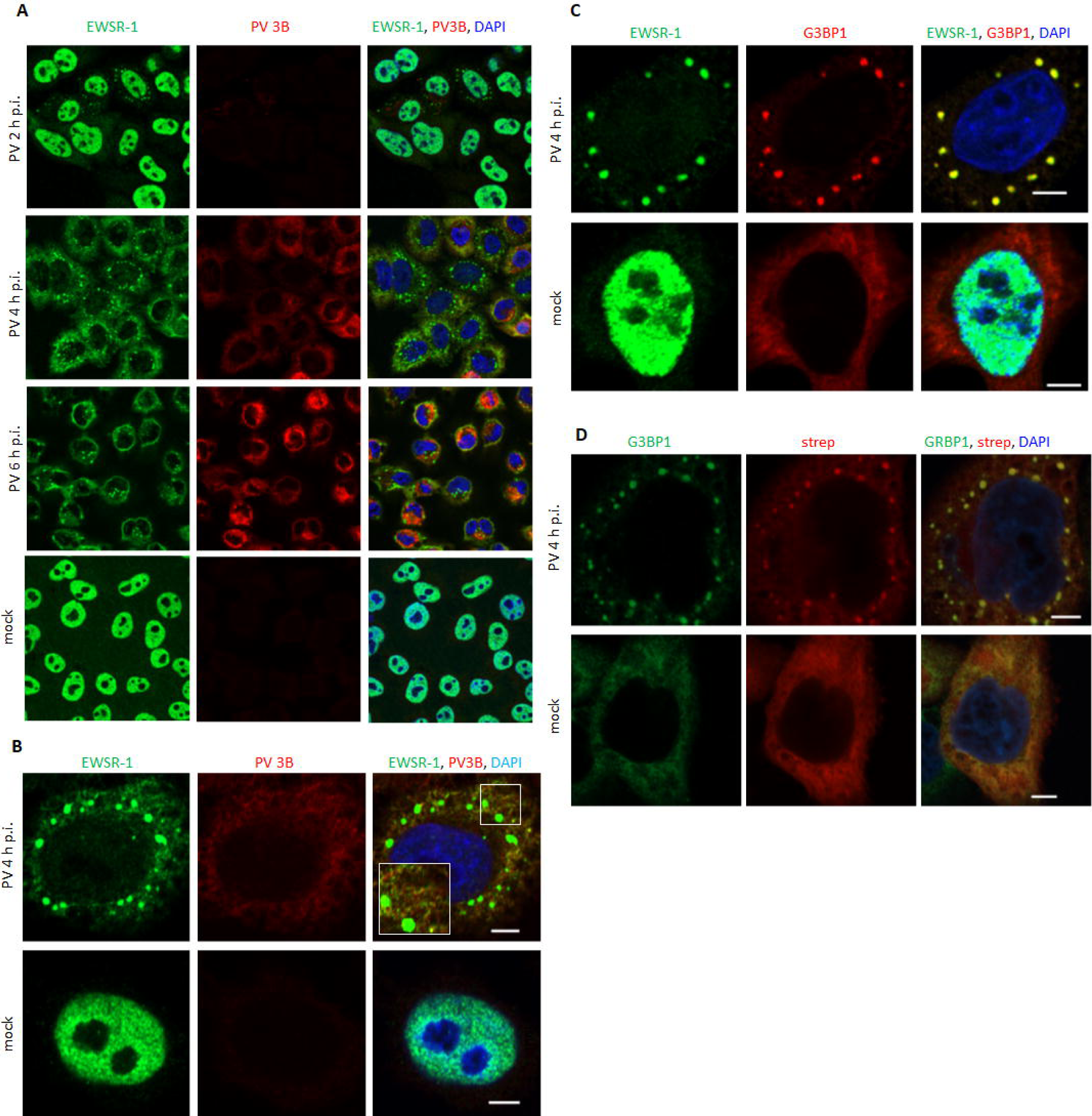
Cytoplasmic translocation of EWSR1 and its association with stress granules upon poliovirus infection. **A.** Confocal images of HeLa cells infected (or mock-infected) with 10 PFU/cell of poliovirus, fixed at 2, 4 and 6 h p.i and stained with antibodies against EWSR1 and the viral replication antigen 3B. **B.** High magnification confocal images of HeLa cells infected (or mock-infected) and processed as in A at 4 h p.i. Note the association of cytoplasmic EWSR1 signal outside of stress granules with the 3B-positive structures. **C.** Confocal images of HeLa cells infected (or mock-infected) as in A, fixed at 4 h.i. and stained with the antibodies against EWSR1 and a stress granule component G3BP1. Scale bar is 10 µm. **D.** Confocal images of HeLa cells stably APEX2-GARG-1060 infected (or mock-infected) as in A, and processed at 4 h p.i. for biotinylation reaction and subsequent staining with antibodies against a stress granule component G3BP1. Scale bar is 5 µm.

Similar to EWSR1, ILF3-90 was localized exclusively in the nuclei of mock-infected cells. Upon poliovirus infection, ILF3-90 signal became exclusively cytoplasmic and was concentrated on the outside margin of the replication organelles, with some of the ILF3-90 distributed within the inner area of the replication organelles (Fig. 8A). The foci of ILF3-90 within the replication organelles strongly colocalized with the dsRNA signal, arguing that its anti-viral activity relies on its dsRNA binding capacity (Fig. 8B).

**Figure 8.**
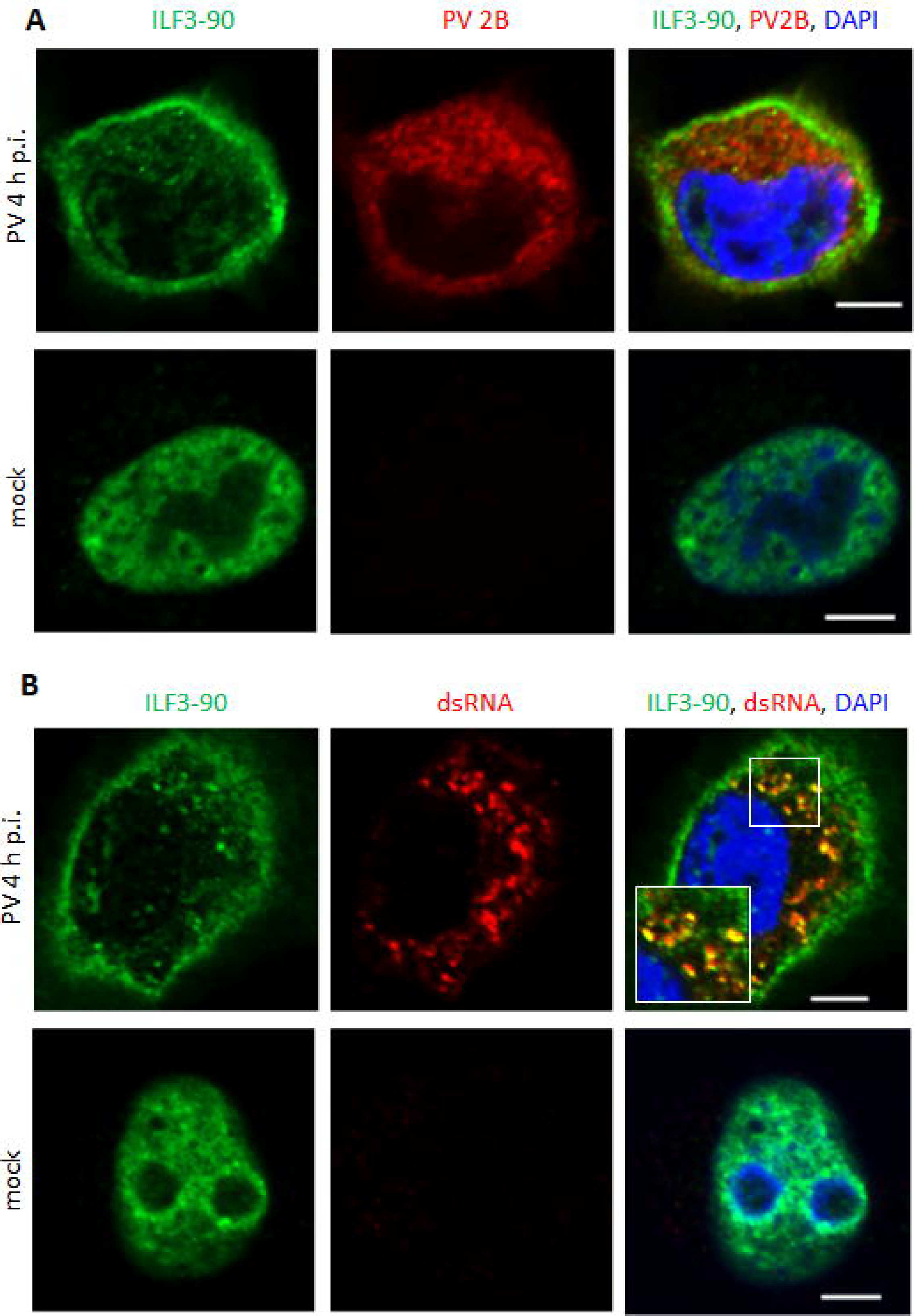
ILF3-90 associates with dsRNA in poliovirus-infected cells. **A, B.** Confocal images of HeLa cells infected (or mock-infected) with 10 PFU/cell of poliovirus, fixed at 4 h p.i. and processed for staining with antibodies against ILF3-90 and a viral antigen 2B, or ILF3-90 and dsRNA, respectively. Scale bar is 10 µm. Note the concentration of ILF3-90 signal on the outer border of the replication organelles and its association with dsRNA inside the replication organelles.

Overall, our data characterize novel proteins affecting the enterovirus replication and demonstrate the complex dynamics of translocation and association with the replication organelles of multiple cellular proteins, underscoring the massive reorganization of the architecture and metabolism of infected cells.

## Discussion

Virtually all stages of the enterovirus replication cycle in a cell are associated with the specialized membranous structures, replication organelles. While their morphological development is extensively documented since the early days of electron microscopy, the landscape of the host and viral proteins on the membranes of the replication organelles and their functional associations are understood only superficially. Here, we used a proximity biotinylation approach to identify proteins localized on the replication organelles in the vicinity of GBF1, a cellular factor indispensable for the RNA replication of all enteroviruses tested so far (47, 61, 106-108). Our GBF1 construct fused to a peroxidase APEX2 had two important modifications. First, the removal of the C-terminal half of GBF1 eliminated the interactions with the cellular proteins non-essential for viral replication (54). Second, our GBF1 construct had a BFA-resistant Sec7 domain, which allowed us to perform infections in the presence of BFA. In these conditions, the endogenous GBF1 was inactivated and the viral replication was supported only by the APEX2-GBF1 fusion increasing the specificity of the detection of proteins relevant for the functioning of the viral replication complexes. The removal of the C-terminal part of GBF1 also resulted in the loss of a specific membrane targeting of the construct in non-infected cells, while upon infection it was effectively recruited to the membranes of the replication organelles. Accordingly, the number of proteins biotinylated by this construct was significantly higher in infected than in non-infected cells.

### Our data in the context of other high throughput studies of cellular factors involved in enterovirus replication

The high throughput methods of identification of host factors involved in the viral replication can be divided into two classes – the ones that are based on a phenotypic signal of depletion of a cellular factor on viral replication, and the unbiased methods that seek to identify all the proteins somehow associated with the viral replication complexes. Each of these approaches can have different technical implementations which have their specific advantages and limitations. The phenotype-based methods usually use siRNA, shRNA or CRISPR-CAS9- based genetic screens targeting the expression of the cellular genes, and by definition would identify functionally important host factors whose depletion either suppresses or promotes the replication. However, these methods may likely miss the proteins important for viral replication that are also essential for cellular viability. Moreover, the prolonged incubation of cells without the expression of a particular protein can induce unpredictable compensatory changes in the expression of other cellular factors which may complicate the interpretation of results. The unbiased approaches, on the other hand, aim to identify all the proteins found at a specific location at a given time, but do not provide immediate information on their functional significance. The spectrum of identified proteins in unbiased approaches depends on the sample enrichment, protein labeling, purification and detection techniques, thus different proteins may be visible or hidden depending on a specific protocol. For example, in our proximity biotinylation approach, we observed that at least one protein (PI4KIIIβ) was found in the biotinylated fraction by western blot but was not identified by subsequent mass-spec analysis. Also, while high enrichment of Arfs on the replication organelles is well documented (82, 83), and at least some amount of Arfs should be close to GBF1, we did not identify Arfs among the proteins biotinylated by our APEX2-GBF1 construct. This may be explained by the intrinsic limitation of the proximity biotinylation which depends on the accessibility of electron-rich amino-acids such as tyrosine, histidine, and tryptophan that can serve as acceptors of biotinphenoxyl radicals (109), i.e. the negative results cannot be unequivocally interpreted as the absence of the protein in the vicinity of the bait. The exposure of suitable amino-acids may be particularly limiting for the detection of small proteins like Arfs.

Previously, both genetic screens and unbiased proteomic approaches were used to characterize host factors involved in the enterovirus replication. RNAi screens were used by Wu et al. to identify factors important for replication of enterovirus A71 in RD cells, by Coyne et al. to find those affecting replication of poliovirus and Coxsackievirus B3 in microvascular endothelial cells, a model of the blood-brain barrier, and by van der Sanden et al. to reveal host proteins restricting or promoting the replication of vaccine strains of poliovirus in Vero cells (110-112). Among the most comprehensive unbiased proteomic screens, van Kuppeveld group used a quantitative characterization of the total protein abundance and phosphorylation status during the time course of Coxsackievirus B3 infection in HeLa cells (113), while Lenarcic and colleagues also used HeLa cells to analyze cellular proteins interacting with poliovirus RNA using labeling of the RNA upon replication with 4-thiouridil followed by crosslinking of RNA-bound proteins (96). Flather et al. characterized the landscape of the nuclear proteins released in the cytoplasm upon rhinovirus infection (27), and Saeed et al. defined the cellular proteins cleaved by enterovirus proteases (114). A comparison of our dataset with those previous reports revealed a particularly strong overlap with the proteins identified as bound to poliovirus RNA (96). From the 81 proteins reported in that study, 38 were also detected by our approach, which supports the localization of GBF1 on the replication organelles close to the RNA replication complexes. Among the genetic screens, our dataset shared 14 genes with those found to be important to support enterovirus 71 replication (112), 13 genes promoting, and six genes restricting poliovirus replication (111). Only VCP (valosine-containing protein, a multifunctional abundant cellular protein involved among other functions in protein quality control at the ER and implicated in the replication of multiple DNA and RNA viruses (115)) and GBF1 were found in our, and two genetic screens (Supplementary Figure 3). Thus, it is important to use different complementary approaches to elucidate the full spectrum of the host proteins involved in viral replication.

### Viral proteins

Among the viral proteins identified in the biotinylated fraction, we observed an increased proportion of the incomplete products of the poliovirus polyprotein cleavage, including some of the fragments that could not be matched with the known intermediates of the polyprotein processing. This suggests that GBF1 on the replication membranes is localized close to the sites of active polyprotein processing. This would be in accordance with the requirement of GBF1 activity for the functioning of the viral RNA replication complexes and the fact that viral replication complexes contain proteins that need to be assembled *in cis*, i.e. derived from the same polyprotein molecule (13, 116-118). On the other hand, an artificial increase of the large polyprotein fragment signal due to a more efficient purification because of a higher proportion of biotinylated amino-acids cannot be excluded. Interestingly, while the large precursors were easily detected in a western blot assay of purified biotinylated proteins, the mass-spectrometry analysis identified viral polypeptides overlapping the 3C-3D but not any other polyprotein cleavage sites, suggesting that most of the polyprotein processing events are very rapid.

### Arfs and Arf effector proteins

GBF1 is an activator of small GTPases of the Arf family and activated Arfs recruit to membranes multiple Arf effector proteins that establish a biochemically distinct local membrane environment (119, 120). The Arf activating function of GBF1 is essential for the enterovirus replication, and all Arf isoforms massively associate with the replication organelles (54, 82, 83). Thus, it is likely that in infected cells Arfs would also recruit the effector proteins to the membranes of the replication organelles which may contribute to the functioning of the replication complexes. Indeed, we identified multiple proteins reported to interact with different Arfs, and the interactors of Arf1 were the most numerous. Given that the depletion of Arf1 affects poliovirus replication much stronger than depletions of other Arfs (83), further detailed investigation of Arf1 effectors on the replication organelles may uncover novel factors important for the viral replication. For our initial characterization of Arf effectors, we chose l-lactate dehydrogenase chains A and B (LDHA and LDHB), reported to interact with Arf1, 4 and 5 (84). However, the siRNA-mediated depletion of LDHA was highly toxic to cells while depletion of LDHB had no effect on the viral replication. Among other Arf effector proteins, we identified in our dataset, Cytohesin2/ARNO, a BFA-insensitive ArfGEF, may be of particular interest. ARNO in non-infected cells regulates membrane trafficking and actin polymerization at the plasma membrane through activation of Arf6, but it can also activate other Arfs, and is itself an Arf1 and Arf6 effector protein (77, 121, 122). Thus, it is likely that in addition to the GBF1- mediated Arf activation, the ARNO-dependent pathway of Arf activation operates on the replication organelles. The existence of such a pathway may contribute to the establishment of resistance to inhibitors of GBF1, such as BFA and similar molecules.

### Both replication-promoting and replication-restricting cellular factors are identified by GBF1-specific biotinylation

We analyzed the effect of depletion of several proteins belonging to different functional groups that were significantly enriched in the dataset from the infected cells on viral replication. AldoA was one of several glycolytic enzymes biotinylated by APEX2- GBF1 in poliovirus-infected cells, and its depletion significantly inhibited the viral replication. The infection-specific biotinylation of multiple glycolytic enzymes suggests an active supply of the replication organelles with the glycolysis pathway-derived metabolites. Recently, the recruitment of glycolytic enzymes to the replication organelles of tombusviruses, a group of positive-strand plant viruses has been discovered, and it was shown that these enzymes are involved in generating a high local level of ATP required to support the viral replication (123-125). Interestingly, the siRNA knockdown of AldoA expression also resulted in a reduction of the cellular ATP level reflected in a lower signal of an ATP-based cell viability test. The AldoA- dependent ATP production was shown to be important for the replication of the Japanese encephalitis virus, a positive-strand RNA virus of the Flaviviridae family (126). AldoA converts fructose 1,6-bisphosphate into glyceraldehyde 3 phosphate (G3P) and dihydroxyacetone phosphate (DHAP) which not only sustain ATP generation in the glycolytic pathway but also serve as substrates for multiple biosynthetic reactions including *de novo* nucleotide synthesis (reviewed in (127)). It is likely that the local nucleotide synthesis sustained in part by the recruitment of the glycolytic enzymes generating the necessary substrates at the replication organelles is a conserved feature of the replication of positive-strand RNA viruses. The importance of *de novo* nucleotide synthesis on the picornavirus replication organelles is highlighted by the previous observations that the partially purified membrane-associated replication complexes more efficiently incorporate in the replicating RNA exogenously added nucleoside mono and diphosphates compared to nucleoside triphosphates (128).

The RNA metabolism proteins were highly enriched in the proteome recovered from the infected cells. Among the RNA-binding proteins, we focused on EWSR1 and ILF3. EWSR1 is an RNA and DNA binding multifunctional protein involved in different networks of regulation of gene expression (129). Depletion of EWSR1 significantly inhibited the replication of both poliovirus and Coxsackie virus B3. The ILF3 gene is expressed as two major isoforms of 110 and 90 KDa which both share the dsRNA binding domain and regulate multiple steps of RNA metabolism in the nucleus and the cytoplasm (130). Since the proteomics data did not distinguish the ILF3 isoforms, we separately targeted the expression of 110K and 90K proteins. Depletion of ILF3-90 but not IF3-110K stimulated the infectious virus yield of poliovirus and Coxsackievirus B3. Thus ILF3-90K may be a broad anti-enterovirus factor. Our data are in contrast with the previously reported specific anti-viral activity of ILF-3 depending on its binding to rhinovirus but not poliovirus IRES (103). This may reflect the difference in the experimental systems used, but the dsRNA-binding capacity of ILF3-90 supported by our data would likely confer the capacity to interfere with the replication of diverse RNA viruses.

Both EWSR1 and ILF3-90 were strictly confined in the nuclei before infection, while in infected cells starting from the middle of the infectious cycle their localization was exclusively cytoplasmic. The disruption of the nucleo-cytoplasmic barrier is caused by the cleavage of nucleoporins by enterovirus protease 2A, which also cleaves a translation initiation factor eIF4G. This results in a rapid inhibition of the mRNA export from the nucleus and cap-dependent translation of cellular mRNAs (18, 20, 21). In addition, the viral proteases 3C and 3CD cleave the core components of RNA polymerase II (131, 132). The rapid and profound inactivation of the cellular gene transcription and translation implies that the proteins present in the cell before infection must contain the full complement of factors required to support the viral replication. This also implies that the cells must dispose of some anti-viral measures ready to be deployed without significant input from the activation of new gene expression. The nuclear depot of RNA-binding proteins thus represents an important resource of both pro- and anti-viral factors. Interestingly, the major ILF3-90 signal was outlining the outer border of the replication organelle area. It is tempting to speculate that this represents the visual manifestation of the protective function of the membranous scaffold of the replication organelles hiding the active replication complexes from the access of anti-viral factors.

Here we selected a few highly abundant proteins for initial characterization, however, the abundance upon the proximity biotinylation-based detection reflects the combination of three variables- the abundance of the protein in a cell, its retention close to the bait, and the exposure of the amino-acids that can accept the biotinphenoxyl radicals. This may skew the representation of the actual enrichment of proteins at the bait construct, thus the less abundant proteins specifically detected at the replication organelles should also be investigated in the future. Overall, our data significantly increased the knowledge of the cellular proteins associated with the enterovirus replication organelles and provide an important resource for the rational approach for the development of antiviral strategies targeting conserved steps of enterovirus replication.

---

**Supplementary Data 1.** Proteomics dataset from LFQ proteomics analysis of the proteins purified upon proximity biotinylation of stable HeLa cell line expressing APEX2-GARG-1060 infected, or mock-infected with 10 PFU/cell of poliovirus at 6 h p.i.

**Supplementary Data 2.** PANTHER Gene Ontology (GO) Overrepresentation Test (Released 20210224) with the proteomics datasets from the poliovirus-infected and mock-infected cells (Supplementary Data 1). GO Ontology database DOI: 10.5281/zenodo.5228828 Released 2021-08-18. Analysis performed: Fisher exact test with Bonferroni correction. Only statistically significant enrichments (p-value <0.05) are shown.

**Supplementary Figure 1.** HeLa cells were transfected with siRNAs specific against the indicated cellular proteins, or non-targeting control siRNA. Polio replicon replication and poliovirus infection assays were performed 72 h post siRNA transfection The total replication signal was calculated as the area under the corresponding kinetics curves. For infection assay, the cells were infected with an MOI of 10 of poliovirus (or mock infected), and processed for western blot with anti-poliovirus 2C antibodies at 4 h p.i. Proteins in red were taken for further analysis. KHDRBS1 (green) is a positive control for a cellular factor known to affect poliovirus replication (86). Each assay was performed at least two time for each protein, repreesnetative results are shown.

Supplementary Table 1. Analysis of potential interactors from the proteomics datasets from the poliovirus-infected and mock-infected cells (Supplementary Data 1) using the Biogrid database of curated interaction data (84, 85).

**Supplementary Table 2.** Analysis of literature on the association of the proteins from the proteomics datasets from the poliovirus-infected and mock-infected cells (Supplementary Data 1) for their involvement in enterovirus replication.

## References

1. Minor P. 2014. The polio endgame. Hum Vaccin Immunother 10:2106–8.

2. Reed Z, Cardosa MJ. 2016. Status of research and development of vaccines for enterovirus 71. Vaccine 34:2967–2970.

3. Oberste MS, Moore D, Anderson B, Pallansch MA, Pevear DC, Collett MS. 2009. In vitro antiviral activity of V-073 against polioviruses. Antimicrob Agents Chemother 53:4501–3.

4. Thibaut HJ, De Palma AM, Neyts J. 2012. Combating enterovirus replication: state-of-the-art on antiviral research. Biochem Pharmacol 83:185–92.

5. Benschop KSM, Van der Avoort HGAM, Duizer E, Koopmans MPG. 2015. Antivirals against enteroviruses: a critical review from a public-health perspective. Antiviral Therapy 20:121–130.

6. Racaniello VR. 2013. Picornaviridae: The Viruses and Their Replication. *In* David M. Knipe PMH (ed), Fields virology, Sixth ed.

7. Lawson MA, Semler BL. 1992. Alternate Poliovirus Nonstructural Protein Processing Cascades Generated by Primary Sites of 3c-Proteinase Cleavage. Virology 191:309–320.

8. Ypma-Wong MF, Semler BL. 1987. Processing determinants required for in vitro cleavage of the poliovirus P1 precursor to capsid proteins. J Virol 61:3181–9.

9. Ypmawong MF, Dewalt PG, Johnson VH, Lamb JG, Semler BL. 1988. Protein 3cd Is the Major Poliovirus Proteinase Responsible for Cleavage of the P1 Capsid Precursor. Virology 166:265–270.

10. Toyoda H, Nicklin MJ, Murray MG, Anderson CW, Dunn JJ, Studier FW, Wimmer E. 1986. A second virus-encoded proteinase involved in proteolytic processing of poliovirus polyprotein. Cell 45:761–70.

11. Giachetti C, Semler BL. 1991. Role of a Viral Membrane Polypeptide in Strand-Specific Initiation of Poliovirus Rna-Synthesis. Journal of Virology 65:2647–2654.

12. Oh HS, Pathak HB, Goodfellow IG, Arnold JJ, Cameron CE. 2009. Insight into Poliovirus Genome Replication and Encapsidation Obtained from Studies of 3B-3C Cleavage Site Mutants. Journal of Virology 83:9370–9387.

13. Egger D, Teterina N, Ehrenfeld E, Bienz K. 2000. Formation of the poliovirus replication complex requires coupled viral translation, vesicle production, and viral RNA synthesis. Journal of Virology 74:6570–6580.

14. Cornell CT, Brunner JE, Semler BL. 2004. Differential rescue of poliovirus RNA replication functions by genetically modified RNA polymerase precursors. Journal of Virology 78:13007–13018.

15. Flather D, Semler BL. 2015. Picornaviruses and nuclear functions: targeting a cellular compartment distinct from the replication site of a positive-strand RNA virus. Front Microbiol 6:594.

16. Baggen J, Thibaut HJ, Strating JRPM, van Kuppeveld FJM. 2018. The life cycle of non-polio enteroviruses and how to target it (vol 16, pg 368, 2018). Nature Reviews Microbiology 16:391–391.

17. Owino CO, Chu JJH. 2019. Recent advances on the role of host factors during non-poliovirus enteroviral infections. Journal of Biomedical Science 26.

18. Park N, Skern T, Gustin KE. 2010. Specific Cleavage of the Nuclear Pore Complex Protein Nup62 by a Viral Protease. Journal of Biological Chemistry 285:28796–28805.

19. Watters K, Inankur B, Gardiner JC, Warrick J, Sherer NM, Yin J, Palmenberg AC. 2017. Differential Disruption of Nucleocytoplasmic Trafficking Pathways by Rhinovirus 2A Proteases. Journal of Virology 91.

20. Gustin KE, Sarnow P. 2001. Effects of poliovirus infection on nucleo-cytoplasmic trafficking and nuclear pore complex composition. Embo Journal 20:240–249.

21. Belov GA, Lidsky PV, Mikitas OV, Egger D, Lukyanov KA, Bienz K, Agol VI. 2004. Bidirectional increase in permeability of nuclear envelope upon poliovirus infection and accompanying alterations of nuclear pores. Journal of Virology 78:10166–10177.

22. Maciejewski S, Nguyen JH, Gomez-Herreros F, Cortes-Ledesma F, Caldecott KW, Semler BL. 2015. Divergent Requirement for a DNA Repair Enzyme during Enterovirus Infections. mBio 7:e01931–15.

23. Virgen-Slane R, Rozovics JM, Fitzgerald KD, Ngo T, Chou W, van Noort GJV, Filippov DV, Gershon PD, Semler BL. 2012. An RNA virus hijacks an incognito function of a DNA repair enzyme. Proceedings of the National Academy of Sciences of the United States of America 109:14634–14639.

24. Hellen CUT, Witherell GW, Schmid M, Shin SH, Pestova TV, Gil A, Wimmer E. 1993. A Cytoplasmic 57-Kda Protein That Is Required for Translation of Picornavirus Rna by Internal Ribosomal Entry Is Identical to the Nuclear Pyrimidine Tract-Binding Protein. Proceedings of the National Academy of Sciences of the United States of America 90:7642–7646.

25. Walter BL, Parsley TB, Ehrenfeld E, Semler BL. 2002. Distinct poly(rC) binding protein KH domain determinants for poliovirus translation initiation and viral RNA replication. Journal of Virology 76:12008–12022.

26. Brunner JE, Nguyen JHC, Roehl HH, Ho TV, Swiderek KM, Semler BL. 2005. Functional interaction of heterogeneous nuclear ribonucleoprotein C with poliovirus RNA synthesis initiation complexes. Journal of Virology 79:3254–3266.

27. Flather D, Nguyen JHC, Semler BL, Gershon PD. 2018. Exploitation of nuclear functions by human rhinovirus, a cytoplasmic RNA virus. Plos Pathogens 14.

28. Flather D, Semler BL. 2015. Picornaviruses and nuclear functions: targeting a cellular compartment distinct from the replication site of a positive-strand RNA virus. Frontiers in Microbiology 6.

29. Holmes AC, Zagnoli-Vieira G, Caldecott KW, Semler BL. 2020. Effects of TDP2/VPg Unlinkase Activity on Picornavirus Infections Downstream of Virus Translation. Viruses-Basel 12.

30. Langereis MA, Feng Q, Nelissen FHT, Virgen-Slane R, van Noort GJV, Maciejewski S, Filippov DV, Semler BL, van Delft FL, van Kuppeveld FJM. 2014. Modification of picornavirus genomic RNA using ‘click’ chemistry shows that unlinking of the VPg peptide is dispensable for translation and replication of the incoming viral RNA. Nucleic Acids Research 42:2473–2482.

31. Jahan N, Wimmer E, Mueller S. 2013. Polypyrimidine Tract Binding Protein-1 (PTB1) Is a Determinant of the Tissue and Host Tropism of a Human Rhinovirus/Poliovirus Chimera PV1(RIPO). Plos One 8.

32. Cheung PKM, Yuan J, Zhang HM, Chau D, Yanagawa B, Suarez A, McManus B, Yang DC. 2005. Specific interactions of mouse organ proteins with the 5’untranslated region of coxsackievirus B3: Potential determinants of viral tissue tropism. Journal of Medical Virology 77:414–424.

33. Viktorova EG, Nchoutmboube JA, Ford-Siltz LA, Iverson E, Belov GA. 2018. Phospholipid synthesis fueled by lipid droplets drives the structural development of poliovirus replication organelles. Plos Pathogens 14.

34. Laufman O, Perrino J, Andino R. 2019. Viral Generated Inter-Organelle Contacts Redirect Lipid Flux for Genome Replication. Cell 178:275-+.

35. Nchoutmboube JA, Viktorova EG, Scott AJ, Ford LA, Pei Z, Watkins PA, Ernst RK, Belov GA. 2013. Increased long chain acyl-Coa synthetase activity and fatty acid import is linked to membrane synthesis for development of picornavirus replication organelles. PLoS Pathog 9:e1003401.

36. Hsu NY, Ilnytska O, Belov G, Santiana M, Chen YH, Takvorian PM, Pau C, van der Schaar H, Kaushik-Basu N, Balla T, Cameron CE, Ehrenfeld E, van Kuppeveld FJM, Altan-Bonnet N. 2010. Viral Reorganization of the Secretory Pathway Generates Distinct Organelles for RNA Replication. Cell 141:799–811.

37. Bauer L, Ferla S, Head SA, Bhat S, Pasunooti KK, Shi WQ, Albulescu L, Liu JO, Brancale A, van Kuppeveld FJM, Strating J. 2018. Structure-activity relationship study of itraconazole, a broad- range inhibitor of picornavirus replication that targets oxysterol-binding protein (OSBP). Antiviral Res 156:55–63.

38. Albulescu L, Bigay J, Biswas B, Weber-Boyvat M, Dorobantu CM, Delang L, van der Schaar HM, Jung YS, Neyts J, Olkkonen VM, van Kuppeveld FJM, Strating J. 2017. Uncovering oxysterol- binding protein (OSBP) as a target of the anti-enteroviral compound TTP-8307. Antiviral Res 140:37–44.

39. Strating JR, van der Linden L, Albulescu L, Bigay J, Arita M, Delang L, Leyssen P, van der Schaar HM, Lanke KH, Thibaut HJ, Ulferts R, Drin G, Schlinck N, Wubbolts RW, Sever N, Head SA, Liu JO, Beachy PA, De Matteis MA, Shair MD, Olkkonen VM, Neyts J, van Kuppeveld FJ. 2015. Itraconazole inhibits enterovirus replication by targeting the oxysterol-binding protein. Cell Rep 10:600–15.

40. Roulin PS, Lotzerich M, Torta F, Tanner LB, van Kuppeveld FJ, Wenk MR, Greber UF. 2014. Rhinovirus uses a phosphatidylinositol 4-phosphate/cholesterol counter-current for the formation of replication compartments at the ER-Golgi interface. Cell Host Microbe 16:677–90.

41. Siltz LAF, Viktorova EG, Zhang B, Kouiavskaia D, Dragunsky E, Chumakov K, Isaacs L, Belov GA. 2014. New Small-Molecule Inhibitors Effectively Blocking Picornavirus Replication. Journal of Virology 88:11091–11107.

42. Alastruey-Izquierdo A, Mellado E, Pelaez T, Peman J, Zapico S, Alvarez M, Rodriguez-Tudela JL, Cuenca-Estrella M, Grp FS. 2013. Phosphatidylinositol 4-Kinase III Beta Is Essential for Replication of Human Rhinovirus and Its Inhibition Causes a Lethal Phenotype In Vivo. Antimicrobial Agents and Chemotherapy 57:3358–3368.

43. Arita M, Kojima H, Nagano T, Okabe T, Wakita T, Shimizu H. 2013. Oxysterol-Binding Protein Family I Is the Target of Minor Enviroxime-Like Compounds. Journal of Virology 87:4252–4260.

44. Arita M, Kojima H, Nagano T, Okabe T, Wakita T, Shimizu H. 2011. Phosphatidylinositol 4-Kinase III Beta Is a Target of Enviroxime-Like Compounds for Antipoliovirus Activity. Journal of Virology 85:2364–2372.

45. Arita M, Takebe Y, Wakita T, Shimizu H. 2010. A bifunctional anti-enterovirus compound that inhibits replication and the early stage of enterovirus 71 infection. Journal of General Virology 91:2734–2744.

46. Lanke KH, van der Schaar HM, Belov GA, Feng Q, Duijsings D, Jackson CL, Ehrenfeld E, van Kuppeveld FJ. 2009. GBF1, a guanine nucleotide exchange factor for Arf, is crucial for coxsackievirus B3 RNA replication. J Virol 83:11940–9.

47. Belov GA, Feng Q, Nikovics K, Jackson CL, Ehrenfeld E. 2008. A Critical Role of a Cellular Membrane Traffic Protein in Poliovirus RNA Replication. Plos Pathogens 4.

48. Wessels E, Duijsings D, Niu TK, Neumann S, Oorschot VM, de Lange F, Lanke KHW, Klumperman J, Henke A, Jackson CL, Melchers WJG, van Kuppeveld FJM. 2006. A viral protein that blocks Arf1-mediated COP-I assembly by inhibiting the guanine nucleotide exchange factor GBF1. Developmental Cell 11:191–201.

49. Ilnytska O, Santiana M, Hsu NY, Du WL, Chen YH, Viktorova EG, Belov G, Brinker A, Storch J, Moore C, Dixon JL, Altan-Bonnet N. 2013. Enteroviruses Harness the Cellular Endocytic Machinery to Remodel the Host Cell Cholesterol Landscape for Effective Viral Replication. Cell Host & Microbe 14:281–293.

50. Lyoo H, van der Schaar HM, Dorobantu CM, Rabouw HH, Strating JRPM, van Kuppeveld FJM. 2019. ACBD3 Is an Essential Pan-enterovirus Host Factor That Mediates the Interaction between Viral 3A Protein and Cellular Protein PI4KB. Mbio 10.

51. Belov GA, Nair V, Hansen BT, Hoyt FH, Fischer ER, Ehrenfeld E. 2012. Complex Dynamic Development of Poliovirus Membranous Replication Complexes. Journal of Virology 86:302–312.

52. Limpens RWAL, van der Schaar HM, Kumar D, Koster AJ, Snijder EJ, van Kuppeveld FJM, Barcena M. 2011. The Transformation of Enterovirus Replication Structures: a Three-Dimensional Study of Single- and Double-Membrane Compartments. Mbio 2.

53. Kaczmarek B, Verbavatz JM, Jackson CL. 2017. GBF1 and Arf1 function in vesicular trafficking, lipid homoeostasis and organelle dynamics. Biology of the Cell 109:391–399.

54. Viktorova EG, Gabaglio S, Meissner JM, Lee E, Moghimi S, Sztul E, Belov GA. 2019. A Redundant Mechanism of Recruitment Underlies the Remarkable Plasticity of the Requirement of Poliovirus Replication for the Cellular ArfGEF GBF1. Journal of Virology 93.

55. Belov GA, Kovtunovych G, Jackson CL, Ehrenfeld E. 2010. Poliovirus replication requires the N- terminus but not the catalytic Sec7 domain of ArfGEF GBF1. Cellular Microbiology 12:1463–1479.

56. Lam SS, Martell JD, Kamer KJ, Deerinck TJ, Ellisman MH, Mootha VK, Ting AY. 2015. Directed evolution of APEX2 for electron microscopy and proximity labeling. Nature Methods 12:51–54.

57. Hung V, Udeshi ND, Lam SS, Loh KH, Cox KJ, Pedram K, Carr SA, Ting AY. 2016. Spatially resolved proteomic mapping in living cells with the engineered peroxidase APEX2. Nature Protocols 11:456–475.

58. Tran JR, Paulson DI, Moresco JJ, Adam SA, Yates JR, Goldman RD, Zheng Y. 2021. An APEX2 proximity ligation method for mapping interactions with the nuclear lamina. J Cell Biol 220.

59. Kärber G. 1931. Beitrag zur kollektiven Behandlung pharmakologischer Reihenversuche. . Archiv f experiment Pathol u Pharmakol:480–483.

60. Viktorova EG, Khattar S, Samal S, Belov GA. 2018. Poliovirus Replicon RNA Generation, Transfection, Packaging, and Quantitation of Replication. Curr Protoc Microbiol 48:15H 4 1–15H 4 15.

61. Lanke KHW, van der Schaar HM, Belov GA, Feng Q, Duijsings D, Jackson CL, Ehrenfeld E, van Kuppeveld FJM. 2009. GBF1, a Guanine Nucleotide Exchange Factor for Arf, Is Crucial for Coxsackievirus B3 RNA Replication. Journal of Virology 83:11940–11949.

62. Pasamontes L, Egger D, Bienz K. 1986. Production of Monoclonal and Monospecific Antibodies against Non-Capsid Proteins of Poliovirus. Journal of General Virology 67:2415–2422.

63. Egger D, Pasamontes L, Bolten R, Boyko V, Bienz K. 1996. Reversible dissociation of the poliovirus replication complex: Functions and interactions of its components in viral RNA synthesis. Journal of Virology 70:8675–8683.

64. Doedens JR, Giddings TH, Kirkegaard K. 1997. Inhibition of endoplasmic reticulum-to-Golgi traffic by poliovirus protein 3A: Genetic and ultrastructural analysis. Journal of Virology 71:9054–9064.

65. Zhang CS, Hawley SA, Zong Y, Li MQ, Wang ZC, Gray A, Ma T, Cui JW, Feng JW, Zhu MJ, Wu YQ, Li TY, Ye ZY, Lin SY, Yin HY, Piao HL, Hardie DGR, Lin SC. 2017. Fructose-1,6-bisphosphate and aldolase mediate glucose sensing by AMPK. Nature 548:112-+.

66. Ahmed NS, Harrell LM, Wieland DR, Lay MA, Thompson VF, Schwartz JC. 2021. Fusion protein EWS-FLI1 is incorporated into a protein granule in cells. Rna 27:920–932.

67. Fei T, Chen YW, Xiao TF, Li W, Cato L, Zhang P, Cotter MB, Bowden M, Lis RT, Zhao SG, Wu Q, Feng FY, Loda M, He HSH, Liu XS, Brown M. 2017. Genome-wide CRISPR screen identifies HNRNPL as a prostate cancer dependency regulating RNA splicing. Proceedings of the National Academy of Sciences of the United States of America 114:E5207–E5215.

68. Fox JT, Shin WK, Caudill MA, Stover PJ. 2009. A UV-responsive Internal Ribosome Entry Site Enhances Serine Hydroxymethyltransferase 1 Expression for DNA Damage Repair. Journal of Biological Chemistry 284:31097–31108.

69. Li L, Yin JY, He FZ, Huang MS, Zhu T, Gao YF, Chen YX, Zhou DB, Chen X, Sun LQ, Zhang W, Zhou HH, Liu ZQ. 2017. Long noncoding RNA&IT SFTA1P&IT promoted apoptosis and increased cisplatin chemosensitivity via regulating the hnRNP-U-GADD45A axis in lung squamous cell carcinoma. Oncotarget 8:97476–97489.

70. Nakamura N, Yamauchi T, Hiramoto M, Yuri M, Naito M, Takeuchi M, Yamanaka K, Kita A, Nakahara T, Kinoyama I, Matsuhisa A, Kaneko N, Koutoku H, Sasamata M, Yokota H, Kawabata S, Furuichi K. 2012. Interleukin Enhancer-binding Factor 3/NF110 Is a Target of YM155, a Suppressant of Survivin. Molecular & Cellular Proteomics 11.

71. Cao D, Haussecker D, Huang Y, Kay MA. 2009. Combined proteomic-RNAi screen for host factors involved in human hepatitis delta virus replication. RNA 15:1971–9.

72. Jin L, Chun J, Pan C, Alesi GN, Li D, Magliocca KR, Kang Y, Chen ZG, Shin DM, Khuri FR, Fan J, Kang S. 2017. Phosphorylation-mediated activation of LDHA promotes cancer cell invasion and tumour metastasis. Oncogene 36:3797–3806.

73. McCleland ML, Adler AS, Shang YL, Hunsaker T, Truong T, Peterson D, Torres E, Li L, Haley B, Stephan JP, Belvin M, Hatzivassiliou G, Blackwood EM, Corson L, Evangelista M, Zha JP, Firestein R. 2012. An Integrated Genomic Screen Identifies LDHB as an Essential Gene for Triple-Negative Breast Cancer. Cancer Research 72:5812–5823.

74. Matsunaga S, Takata H, Morimoto A, Hayashihara K, Higashi T, Akatsuchi K, Mizusawa E, Yamakawa M, Ashida M, Matsunaga TM, Azuma T, Uchiyama S, Fukui K. 2012. RBMX: A Regulator for Maintenance and Centromeric Protection of Sister Chromatid Cohesion. Cell Reports 1:299–308.

75. Kall L, Storey JD, Noble WS. 2008. Non-parametric estimation of posterior error probabilities associated with peptides identified by tandem mass spectrometry. Bioinformatics 24:I42–I48.

76. Mi HY, Ebert D, Muruganujan A, Mills C, Albou LP, Mushayamaha T, Thomas PD. 2021. PANTHER version 16: a revised family classification, tree-based classification tool, enhancer regions and extensive API. Nucleic Acids Research 49:D394–D403.

77. Jackson CL, Casanova JE. 2000. Turning on ARF: the Sec7 family of guanine-nucleotide-exchange factors. Trends in Cell Biology 10:60–67.

78. Bhatt JM, Hancock W, Meissner JM, Kaczmarczyk A, Lee E, Viktorova E, Ramanadham S, Belov GA, Sztul E. 2019. Promiscuity of the catalytic Sec7 domain within the guanine nucleotide exchange factor GBF1 in ARF activation, Golgi homeostasis, and effector recruitment. Molecular Biology of the Cell 30:1523–1535.

79. Pocognoni CA, Viktorova EG, Wright J, Meissner JM, Sager G, Lee E, Belov GA, Sztul E. 2018. Highly conserved motifs within the large Sec7 ARF guanine nucleotide exchange factor GBF1 target it to the Golgi and are critical for GBF1 activity. American Journal of Physiology-Cell Physiology 314:C675–C689.

80. Wright J, Kahn RA, Sztul E. 2014. Regulating the large Sec7 ARF guanine nucleotide exchange factors: the when, where and how of activation. Cellular and Molecular Life Sciences 71:3419–3438.

81. Olson MG, Widner RE, Jorgenson LM, Lawrence A, Lagundzin D, Woods NT, Ouellette SP, Rucks EA. 2019. Proximity Labeling To Map Host-Pathogen Interactions at the Membrane of a Bacterium-Containing Vacuole in Chlamydia trachomatis-Infected Human Cells. Infection and Immunity 87.

82. Belov GA, Altan-Bonnet N, Kovtunovych G, Jackson CL, Lippincott-Schwartz J, Ehrenfeld E. 2007. Hijacking components of the cellular secretory pathway for replication of poliovirus RNA. Journal of Virology 81:558–567.

83. Moghimi S, Viktorova E, Zimina A, Szul T, Sztul E, Belov GA. 2021. Enterovirus Infection Induces Massive Recruitment of All Isoforms of Small Cellular Arf GTPases to the Replication Organelles. Journal of Virology 95.

84. Oughtred R, Rust J, Chang C, Breitkreutz BJ, Stark C, Willems A, Boucher L, Leung G, Kolas N, Zhang F, Dolma S, Coulombe-Huntington J, Chatr-aryamontri A, Dolinski K, Tyers M. 2021. The BioGRID database: A comprehensive biomedical resource of curated protein, genetic, and chemical interactions. Protein Science 30:187–200.

85. Stark C, Breitkreutz BJ, Reguly T, Boucher L, Breitkreutz A, Tyers M. 2006. BioGRID: a general repository for interaction datasets. Nucleic Acids Research 34:D535–D539.

86. McBride AE, Schlegel A, Kirkegaard K. 1996. Human protein Sam68 relocalization and interaction with poliovirus RNA polymerase in infected cells. Proceedings of the National Academy of Sciences of the United States of America 93:2296–2301.

87. Herold J, Andino R. 2001. Poliovirus RNA replication requires genome circularization through a protein-protein bridge. Molecular Cell 7:581–591.

88. Toyoda H, Franco D, Fujita K, Paul AV, Wimmer E. 2007. Replication of poliovirus requires binding of the poly(rC) binding protein to the cloverleaf as well as to the adjacent C-rich spacer sequence between the cloverleaf and the internal ribosomal entry site. Journal of Virology 81:10017–10028.

89. Blyn LB, Towner JS, Semler BL, Ehrenfeld E. 1997. Requirement of Poly(rC) binding protein 2 for translation of poliovirus RNA. Journal of Virology 71:6243–6246.

90. Parsley TB, Towner JS, Blyn LB, Ehrenfeld E, Semler BL. 1997. Poly (rC) binding protein 2 forms a ternary complex with the 5’-terminal sequences of poliovirus RNA and the viral 3CD proteinase. Rna 3:1124–1134.

91. Kafasla P, Lin H, Curry S, Jackson RJ. 2011. Activation of picornaviral IRESs by PTB shows differential dependence on each PTB RNA-binding domain. Rna 17:1120–1131.

92. Kafasla P, Morgner N, Robinson CV, Jackson RJ. 2010. Polypyrimidine tract-binding protein stimulates the poliovirus IRES by modulating eIF4G binding. Embo Journal 29:3710–3722.

93. Pedley AM, Benkovic SJ. 2017. A New View into the Regulation of Purine Metabolism: The Purinosome. Trends in Biochemical Sciences 42:141–154.

94. Wang YJ, Wang WS, Xu L, Zhou XY, Shokrollahi E, Felczak K, van der Laan LJW, Pankiewicz KW, Sprengers D, Raat NJH, Metselaar HJ, Peppelenbosch MP, Pan QW. 2016. Cross Talk between Nucleotide Synthesis Pathways with Cellular Immunity in Constraining Hepatitis E Virus Replication. Antimicrobial Agents and Chemotherapy 60:2834–2848.

95. Ariav Y, Ch’ng JH, Christofk HR, Ron-Harel N, Erez A. 2021. Targeting nucleotide metabolism as the nexus of viral infections, cancer, and the immune response. Science Advances 7.

96. Lenarcic EM, Landry DM, Greco TM, Cristea IM, Thompson SR. 2013. Thiouracil Cross-Linking Mass Spectrometry: a Cell-Based Method To Identify Host Factors Involved in Viral Amplification. Journal of Virology 87:8697–8712.

97. Watson SF, Bellora N, Macias S. 2020. ILF3 contributes to the establishment of the antiviral type I interferon program. Nucleic Acids Research 48:116–129.

98. Li X, Liu CX, Xue W, Zhang Y, Jiang S, Yin QF, Wei J, Yao RW, Yang L, Chen LL. 2017. Coordinated circRNA Biogenesis and Function with NF90/NF110 in Viral Infection. Molecular Cell 67:214-+.

99. Gomila RC, Martin GW, Gehrke L. 2011. NF90 Binds the Dengue Virus RNA 3 ’ Terminus and Is a Positive Regulator of Dengue Virus Replication. Plos One 6.

100. Isken O, Baroth M, Grassmann CW, Weinlich S, Ostareck DH, Ostareck-Lederer A, Behrens SE. 2007. Nuclear factors are involved in hepatitis C virus RNA replication. Rna 13:1675–1692.

101. Reichman TW, Muniz LC, Mathews MB. 2002. The RNA binding protein nuclear factor 90 functions as both a positive and negative regulator of gene expression in mammalian cells. Molecular and Cellular Biology 22:343–356.

102. Patino C, Haenni AL, Urcuqui-Inchima S. 2015. NF90 isoforms, a new family of cellular proteins involved in viral replication? Biochimie 108:20–24.

103. Merrill MK, Gromeier M. 2006. The double-stranded RNA binding protein 76 : NF45 heterodimer inhibits translation initiation at the rhinovirus type 2 internal ribosome entry site. Journal of Virology 80:6936–6942.

104. Mamczur P, Gamian A, Kolodziej J, Dziegiel P, Rakus D. 2013. Nuclear localization of aldolase A correlates with cell proliferation. Biochimica Et Biophysica Acta-Molecular Cell Research 1833:2812–2822.

105. Dougherty JD, Tsai WC, Lloyd RE. 2015. Multiple Poliovirus Proteins Repress Cytoplasmic RNA Granules. Viruses-Basel 7:6127–6140.

106. Farhat R, Ankavay M, Lebsir N, Gouttenoire J, Jackson CL, Wychowski C, Moradpour D, Dubuisson J, Rouille Y, Cocquerel L. 2018. Identification of GBF1 as a cellular factor required for hepatitis E virus RNA replication. Cellular Microbiology 20.

107. Verheije MH, Raaben M, Mari M, Lintelo EGT, Reggiori F, van Kuppeveld FJM, Rottier PJM, de Haan CAM. 2008. Mouse hepatitis coronavirus RNA replication depends on GBF1-mediated ARF1 activation. Plos Pathogens 4.

108. Goueslain L, Alsaleh K, Horellou P, Roingeard P, Descamps V, Duverlie G, Ciczora Y, Wychowski C, Dubuisson J, Rouille Y. 2010. Identification of GBF1 as a Cellular Factor Required for Hepatitis C Virus RNA Replication. Journal of Virology 84:773–787.

109. Rhee HW, Zou P, Udeshi ND, Martell JD, Mootha VK, Carr SA, Ting AY. 2013. Proteomic Mapping of Mitochondria in Living Cells via Spatially Restricted Enzymatic Tagging. Science 339:1328–1331.

110. Coyne CB, Bozym R, Morosky SA, Hanna SL, Mukherjee A, Tudor M, Kim KS, Cherry S. 2011. Comparative RNAi Screening Reveals Host Factors Involved in Enterovirus Infection of Polarized Endothelial Monolayers. Cell Host & Microbe 9:70–82.

111. van der Sanden SMG, Wu WL, Dybdahl-Sissoko N, Weldon WC, Brooks P, O’Donnell J, Jones LP, Brown C, Tompkins SM, Oberste MS, Karpilow J, Tripp RA. 2016. Engineering Enhanced Vaccine Cell Lines To Eradicate Vaccine-Preventable Diseases: the Polio End Game. Journal of Virology 90:1694–1704.

112. Wu KX, Phuektes P, Kumar P, Goh GYL, Moreau D, Chow VTK, Bard F, Chu JJH. 2016. Human genome-wide RNAi screen reveals host factors required for enterovirus 71 replication. Nature Communications 7.

113. Giansanti P, Strating JRPM, Defourny KAY, Cesonyte I, Bottino AMS, Post H, Viktorova EG, Ho VQT, Langereis MA, Belov GA, Nolte-t Hoen ENM, Heck AJR, van Kuppeveld FJM. 2020. Dynamic remodelling of the human host cell proteome and phosphoproteome upon enterovirus infection. Nature Communications 11.

114. Saeed M, Kapell S, Hertz NT, Wu XF, Bell K, Ashbrook AW, Mark MT, Zebroski HA, Neal ML, Flodstrom-Tullberg M, MacDonald MR, Aitchison JD, Molina H, Rice CM. 2020. Defining the proteolytic landscape during enterovirus infection. Plos Pathogens 16.

115. Das P, Dudley JP. 2021. How Viruses Use the VCP/p97 ATPase Molecular Machine. Viruses-Basel 13.

116. Teterina NL, Zhou WD, Cho MW, Ehrenfeld E. 1995. Inefficient Complementation Activity of Poliovirus 2c and 3d Proteins for Rescue of Lethal Mutations. Journal of Virology 69:4245–4254.

117. Towner JS, Mazanet MM, Semler BL. 1998. Rescue of defective poliovirus RNA replication by 3AB-containing precursor polyproteins. Journal of Virology 72:7191–7200.

118. Pathak HB, Oh HS, Goodfellow IG, Arnold JJ, Cameron CE. 2008. Picornavirus genome replication: roles of precursor proteins and rate-limiting steps in oriI-dependent VPg uridylylation. J Biol Chem 283:30677–88.

119. Nie ZZ, Hirsch DS, Randazzo PA. 2003. Arf and its many interactors. Current Opinion in Cell Biology 15:396–404.

120. Jackson CL. 2014. Arf Proteins and Their Regulators: At the Interface Between Membrane Lipids and the Protein Trafficking Machinery. . *In* A. W (ed), Ras Superfamily Small G Proteins: Biology and Mechanisms 2 doi:10.1007/978-3-319-07761-1_8. Springer, Cham.

121. Cohen LA, Honda A, Varnai P, Brown FD, Balla T, Donaldson JG. 2007. Active Arf6 recruits ARNO/cytohesin GEFs to the PM by binding their PH domain. Molecular Biology of the Cell 18:2244–2253.

122. Li HS, Shome K, Rojas R, Rizzo MA, Vasudevan C, Fluharty E, Santy LC, Casanova JE, Romero G. 2003. The Guanine Nucleotide Exchange Factor ARNO mediates the activation of ARF and phospholipase D by insulin. Bmc Cell Biology 4:1–10.

123. Prasanth KR, Chuang CK, Nagy PD. 2017. Co-opting ATP-generating glycolytic enzyme PGK1 phosphoglycerate kinase facilitates the assembly of viral replicase complexes. Plos Pathogens 13.

124. Chuang CK, Prasanth KR, Nagy PD. 2017. The Glycolytic Pyruvate Kinase Is Recruited Directly into the Viral Replicase Complex to Generate ATP for RNA Synthesis. Cell Host & Microbe 22:639-+.

125. Lin WW, Liu YY, Molho M, Zhang SJ, Wang LS, Xie LH, Nagy PD. 2019. Co-opting the fermentation pathway for tombusvirus replication: Compartmentalization of cellular metabolic pathways for rapid ATP generation. Plos Pathogens 15.

126. Tien CF, Cheng SC, Ho YP, Chen YS, Hsu JH, Chang RY. 2014. Inhibition of aldolase A blocks biogenesis of ATP and attenuates Japanese encephalitis virus production. Biochemical and Biophysical Research Communications 443:464–469.

127. Chang YC, Yang YC, Tien CP, Yang CJ, Hsiao M. 2018. Roles of Aldolase Family Genes in Human Cancers and Diseases. Trends in Endocrinology and Metabolism 29:549–559.

128. Koonin EV, Agol VI. 1984. Encephalomyocarditis Virus-Replication Complexes Preferentially Utilizing Nucleoside Diphosphates as Substrates for Viral-Rna Synthesis - Nucleotide Kinases Specifically Associated with the Complex Channel Rna Precursor. European Journal of Biochemistry 144:249–254.

129. Lee J, Nguyen PT, Shim HS, Hyeon SJ, Im H, Choi MH, Chung S, Kowall NW, Lee SB, Ryu H. 2019. EWSR1, a multifunctional protein, regulates cellular function and aging via genetic and epigenetic pathways. Biochimica Et Biophysica Acta-Molecular Basis of Disease 1865:1938–1945.

130. Castella S, Bernard R, Corno M, Fradin A, Larcher JC. 2015. Ilf3 and NF90 functions in RNA biology. Wiley Interdisciplinary Reviews-Rna 6:243–256.

131. Kundu P, Raychaudhuri S, Tsai W, Dasgupta A. 2005. Shutoff of RNA polymerase II transcription by poliovirus involves 3C protease-mediated cleavage of the TATA-binding protein at an alternative site: Incomplete shutoff of transcription interferes with efficient viral replication. Journal of Virology 79:9702–9713.

132. Sharma R, Raychaudhuri S, Dasgupta A. 2004. Nuclear entry of poliovirus protease-polymerase precursor 3CD: implications for host cell transcription shut-off. Virology 320:195–205.

