## Supplementary figures and images for "A Proximity Biotinylation Assay with a Host Protein Bait Reveals Multiple Factors Modulating Enterovirus Replication"

### Supplementary Figure 1

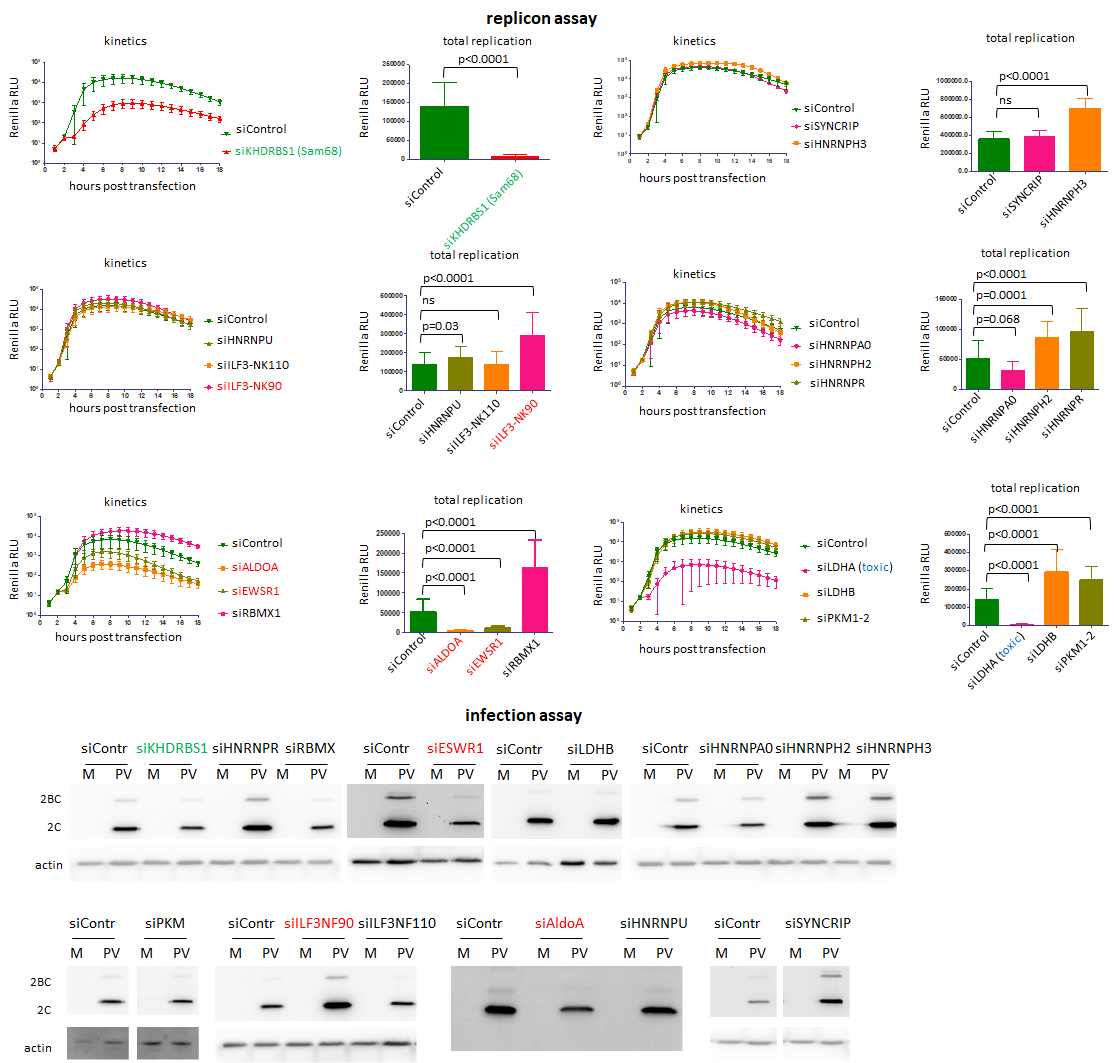
